# PhysioMap: an ontology-grounded causal knowledge graph of human physiology

**DOI:** 10.64898/2026.08.03.742471

**Authors:** Robert Hoehndorf, Paul N. Schofield, Georgios V. Gkoutos

## Abstract

Computational physiology needs representations that connect traits across biological scales while distinguishing causal, constitutive, and mathematical dependencies. We present PhysioMap, an ontology-grounded knowledge base of contextualized physiological traits and precisely defined relation types. A versioned projection maps entailed ontology patterns to a typed causal knowledge graph that constrains quantitative structural causal models. Derivative signs provide a separate qualitative abstraction, which the PhysioMap solver uses to analyze steady-state responses in the presence of feedback. A stratified expert review across all relation types supported most sampled relations and isolated a minority for correction or further investigation. In a rare metabolic disease application, nearly all determinate predictions agreed with the HPO-derived reference before post-hoc review; after the discordant reference directions were excluded, all remaining determinate predictions agreed. Shortest signed paths produced directional errors, particularly on cases for which the PhysioMap solver did not determine a direction, indicating that its abstentions concentrated difficult cases. Abduction usually narrowed the candidate set but often did not identify a unique cause. PhysioMap therefore connects ontology-grounded physiological content to interventional prediction and abduction under incomplete quantitative knowledge. Because PhysioMap curation and the HPO-derived reference may share supporting literature, and because abduction used a closed candidate pool, these analyses do not constitute independent clinical validation.

## 1. Introduction

Computational physiology must represent processes, participants, and traits in specified biological contexts. Here, we define a physiological trait as a quality borne by an entity or process in such a context. A physiological representation should support prediction, which computes the consequences of an intervention, and abduction, which identifies interventions that could account for observations. It must also connect molecular, cellular, tissue, organ, and whole-body scales while preserving feedback. These operations require traits to denote endogenous random variables connected through mechanisms that admit intervention, even when complete equations are unavailable.

Rare metabolic diseases provide a concrete application. A pathogenic variant may alter an enzyme or transporter, metabolic flux, and a measurable endophenotype, while homeostatic control can amplify, attenuate, or oppose the effect. Forward prediction asks which contextualized variables rise or fall after the system reaches a new state following an intervention on a lesion variable. A sound method should abstain when available knowledge does not determine a direction. The same query can represent drug-target activation or inhibition; abduction instead asks which intervention best explains observed traits.

Directed molecular resources such as SIGNOR, OmniPath, INDRA, Reactome, and KEGG encode regulatory, signaling, and reaction knowledge.^1–5^ Network-propagation methods spread evidence beyond direct annotations,^6,7^ while sign-consistency or optimization methods constrain explanations to observed molecular changes.^8,9^ These approaches provide broad coverage, but local traversal does not specify the response of a coupled system to intervention. Converging paths may reinforce or oppose one another, and feedback can leave the net response unresolved.

Constraint-based metabolic reconstructions predict biomarkers of enzyme defects, while metabolomic propagation ranks disease genes.^10,11^ Parameterized differential equations represent feedback explicitly, and CellML, BioModels, HumMod, and Physiome make quantitative subsystem models reusable.^12–15^ Multiscale integration, however, requires compatible variables, equations, and parameters that are often unavailable. Earlier physio-maps and Chalkboard instead combined ontology-grounded physiological processes with qualitative inference;^16,17^ the Physiome Project and NIH Whole Person Initiative seek reusable multiorgan representations.^12,18,19^

Biomedical ontologies formalize processes, anatomy, cells, chemicals, and proteins.^20–24^ Entity–quality patterns combine PATO qualities with these classes to define traits and phenotypes that correspond to physiological variables.^25–27^ OWL reasoning classifies these traits and propagates logical relations such as parthood. OWL does not, by itself, give “may cause” statements a standard instance-level semantics,^28^ nor does a directed ontology relation define the response of a system to an intervention.

In a structural causal model (SCM), a perfect intervention sets a target variable to a specified value, replacing the equation that normally determines it while leaving all other equations unchanged.^29,30^ Physiological feedback requires causal semantics that accommodate cycles, which violate the acyclicity assumed by many graphical models.^31,32^ The remaining problem is to connect this interventional semantics to ontological knowledge without conflating causal influence with parthood, production, quantitative identity, or other typed dependencies. Ontology entailments determine which traits and relations enter a model, while the SCM semantics determines how an intervention propagates through them.

We developed PhysioMap to make this connection. We collected physiological statements from textbooks, systems biology models, curated resources, and direct expert curation; classified them by relation type; and encoded them as patterns in an OWL knowledge base that grounds physiological traits in external ontologies. A versioned projection transforms its deductive closure into a causal knowledge graph (CKG), a knowledge graph with explicitly interpreted causal edges.^33^ Release v1.1.1 represents 1,699 traits and 2,387 typed relations across five relation types.

In the SCM semantics, projected traits denote quantitative random variables, and the five relation types constrain structural functions, identities, constitutive maps, or mixed derivatives. Derivative signs form a separate abstraction of these constraints. We implement the PhysioMap solver, a feedback-aware algorithm for the first-order derivative-sign abstraction, and a separate qualitative modulation query. We evaluate the PhysioMap solver through forward prediction of directional phenotypes after interventions on gene-associated lesion variables and through abduction of candidate lesion interventions from their predicted phenotype profiles.

### Data and code availability

The OWL knowledge base, projection registry, causal model, release metadata, evaluation code, and archived expert review are available at https://github.com/bio-ontology-research-group/physiomap/. The supplementary material is available directly at https://github.com/bio-ontology-research-group/physiomap/releases/download/v1.1.1/supplement.pdf. An interactive demo for browsing the typed map and exploring qualitative intervention predictions is available at https://bio2vec.net/physiomap/. Code uses BSD 3-Clause; the map, benchmark data, and documentation use CC BY 4.0.

## 2. Methods

### 2.1. Ontology and trait semantics

We first represent PhysioMap as an OWL 2 terminological knowledge base (TBox) *K* with standard OWL semantics.^34^ Its upper level distinguishes continuants, processes, and traits. Imported modules provide classes for chemicals, proteins, cells, anatomy, processes, and qualities from ChEBI, PR, CL, Uberon, GO, and PATO.^20–25^

We use the entity–quality phenotype pattern to define a trait as a quality borne by an entity or process in a biological context.^26,27^ A named trait *T*_*i*_ has the necessary conditions

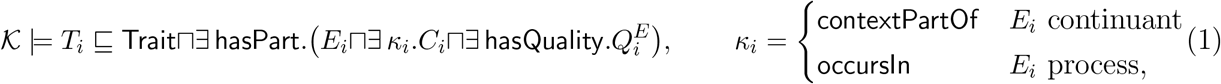

where *E*_*i*_ is the characterized entity or process, *C*_*i*_ its recorded context, and 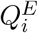 an entity-specific subclass of the PATO quality *Q*_*i*_. We omit the context conjunct when no context is recorded. The relation `contextPartOf` localizes a continuant, whereas `occursIn` localizes a process and propagates along parthood (`occursIn º partOf ⊑ occursIn`). The PATO hierarchy determines whether a quality requires a continuant or process bearer.

Parthood also connects traits across scales. For a quality that composes over parts, the TBox defines *G*_*E,Q*_ ≡ ∃ `hasPart`.(Ê ⊓∃ `hasQuality`.*Q*) with Ê the reflexive `partOf` closure of *E*. ELK can therefore classify plasma volume, erythrocyte volume, and blood volume under *volume of blood or its parts*. This entailment records a shared bearer context; it neither sums part volumes nor makes plasma volume a subtype of blood volume. We create these grouping classes only for composable qualities. For example, volume is extensive and composes over parts, whereas pressure is intensive and does not.

We also encode relations between traits as OWL axioms. A causal axiom states a type-level capacity, not that every instance of the target trait is caused by an instance of the source. The subclass axiom *T*_*t*_ ⊑ ∃ `causedBy`.*T*_*s*_ would make the latter, stronger claim. A fully dispositional formulation, *T*_*t*_ ⊑ ∃ `hasDisposition`.(∀ realizedBy.∃ `hasParticipant`.*T*_*s*_), mixes existential and universal restrictions and leaves OWL 2 EL. We therefore use the collection pattern developed for biological function.^35^ Each trait *T* has an associated collection class *T* ^all^, with *T* ⊑ ∃ memberOf.*T* ^all^ and *T* ^all^ ⊑ ∃ `hasMember`.*T*. A causal axiom then states that some member of the target collection is caused by an instance of the source, 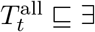 hasMember.(∃ causedBy.*T*_*s*_). These collection-pattern axioms are existential and remain in OWL 2 EL. The DL artifact additionally states that the collection is homogeneous and has one instance, using *T* ^all^ ⊑ ∀ hasMember.*T*, which lies outside EL, and a singleton axiom. Neither is used for classification. Production and modulation are type-level mechanism claims of the same kind and take the same form with `producedBy` and `modulates`, the latter qualifying the causal axiom it changes. These axioms give PhysioMap its logical structure; the projection below maps that structure to variables and supplies the interventional semantics.

### 2.2. Projection from ontology to a causal knowledge graph

We next project the OWL knowledge base into a causal knowledge graph (CKG), that is, a knowledge graph with an explicitly interpreted causal edge type.^33^ Let *V* be its named trait classes and *E*_*c*_ ⊆ *V ×V* its causal-influence edges. The CKG also contains the typed production,

constitution, quantitative-identity, and modulation relation instances defined below. A binary relation instance *T*_*s*_ → *T*_*t*_ labeled *r* is present whenever *K* entails a relational pattern that the finite, versioned registry Π registers for relation type *r*. Each pattern *ρ* ∈ Π defines an arity, direction, relation type, OWL class-expression premise φ_*ρ*_, and the relation instance produced when that premise is entailed:

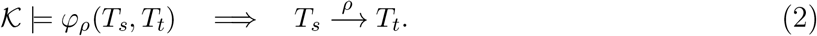

Higher-arity patterns analogously produce typed hyperedges. The causal pattern, for example, has 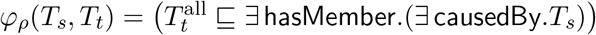, so the entailment holding of ACE activity and angiotensin II yields the causal edge between those two traits.

Because the projection operates on entailments, the CKG depends on what the reasoner derives rather than on the syntactic form of an axiom.^36,37^

### 2.3. Structural causal model semantics for typed relations

The projection targets a class M(*K*, Π) of quantitative structural causal models, not a signed graph. Trait *T*_*i*_ denotes an endogenous random variable *X*_*i*_, and *x*_*i*_ a realized value. Exogenous random variables *U*_*i*_ enter the differentiable dynamics 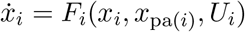. We assume a locally unique equilibrium *x*^*^ with a Hurwitz Jacobian and local self-regulation 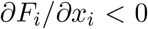 A perfect intervention do(*x*_*A*_ = *θ*_*A*_) replaces the mechanism or defining equation of each *x*_*a*_ by *x*_*a*_ = *θ*_*a*_, removes its incoming functional dependencies, and leaves all other equations unchanged.^29,38,39^ Relation typing matters because it determines whether an intervention replaces a causal mechanism, a constitutive constraint, or a quantitative identity.

To define a direct effect, we let *x*_*t*_ re-equilibrate after an intervention on *x*_*s*_ while keeping every other variable at its reference value. Suppl. Sec. S6.1 proves the sign identity:

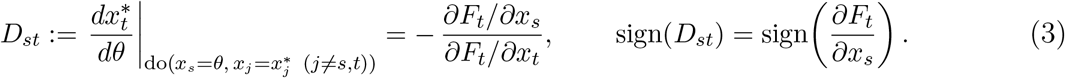

For a multivariate function, *σ*_*j*_ denotes the sign of its partial derivative with respect to argument *j*. Table 1 gives the quantitative constraint for each first-order relation type, its derivative-sign abstraction, and a PhysioMap example.

**Table 1.**
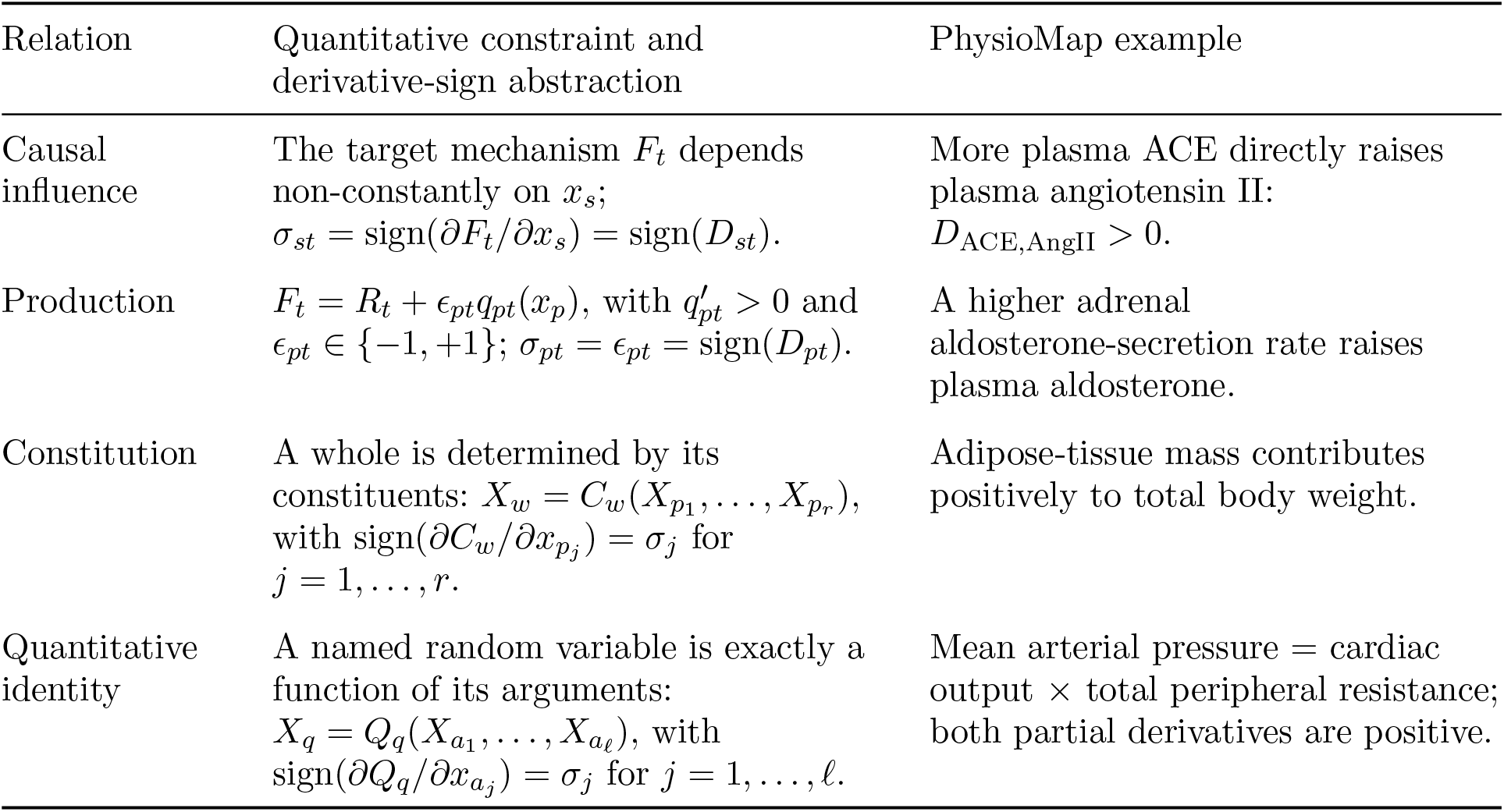
Structural causal model semantics, derivative-sign abstractions, and examples for the first-order relation types. *D*_*st*_ is the direct effect in Eq. (3).

| Relation | Quantitative constraint and derivative-sign abstraction | PhysioMap example |
| --- | --- | --- |
| Causal influence | The target mechanism $F_t$ depends non-constantly on $x_s$;<br>$\sigma_{st} = \text{sign}(\partial F_t / \partial x_s) = \text{sign}(D_{st})$. | More plasma ACE directly raises plasma angiotensin II:<br>$D_{\text{ACE}, \text{AngII}} > 0$. |
| Production | $F_t = R_t + \epsilon_{pt} q_{pt}(x_p)$, with $q'_{pt} > 0$ and $\epsilon_{pt} \in \{-1, +1\}$; $\sigma_{pt} = \epsilon_{pt} = \text{sign}(D_{pt})$. | A higher adrenal aldosterone-secretion rate raises plasma aldosterone. |
| Constitution | A whole is determined by its constituents: $X_w = C_w(X_{p_1}, \dots, X_{p_r})$, with $\text{sign}(\partial C_w / \partial x_{p_j}) = \sigma_j$ for $j = 1, \dots, r$. | Adipose-tissue mass contributes positively to total body weight. |
| Quantitative identity | A named random variable is exactly a function of its arguments:<br>$X_q = Q_q(X_{a_1}, \dots, X_{a_\ell})$, with $\text{sign}(\partial Q_q / \partial x_{a_j}) = \sigma_j$ for $j = 1, \dots, \ell$. | Mean arterial pressure = cardiac output × total peripheral resistance; both partial derivatives are positive. |

Sign annotations belong to a derived abstraction of the quantitative constraints. The abstract value *σ* = `?` requires non-constant direct dependence but places no sign constraint on *D*_*st*_; it is not a value of *X*_*s*_, *X*_*t*_, *x*_*s*_, or *x*_*t*_. Constitution and quantitative identity are directed: an intervention on the result replaces its defining equation rather than propagating backward to its arguments.

A modulation (*m*; (*s, t*), *µ*), where *µ* is the mixed-derivative sign, is a (node, causal-influence edge) tuple in *H* ⊆ *V × E*_*c*_. Its semantics is a paired intervention on modulator and source,

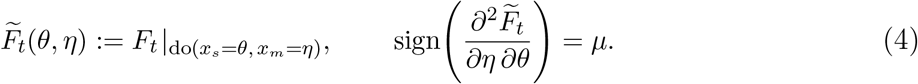

With other arguments held fixed, the sign states whether do(*x*_*m*_) makes the direct effect of do(*x*_*s*_) more or less positive. For example, higher cortisol makes a fixed increase in norepinephrine produce a larger increase in total peripheral resistance. The current projection retains modulation in *H*, which the first-order solver does not read. It neither derives the required parent *m* → *t* from a modulation nor changes the endpoint direction it returns. A first-order effect of the modulator must therefore be authored separately as a causal influence, which remains an ordinary edge in *E*_*c*_. A separate qualitative query combines a modulator’s first-order response with the mixed-derivative sign to report whether the signed slope becomes more or less positive. Suppl. Secs. S9.2 and S9.3 give the full second-order and population interpretations.

### 2.4. First-order derivative-sign abstraction and feedback-aware prediction

The quantitative SCM supplies the semantics; the PhysioMap solver reads only its first-order derivative-sign abstraction. The direct effects above keep the rest of the system fixed. Forward prediction instead asks for the total effect after every other variable re-equilibrates. At the assumed equilibrium *F* (*x*^*^, *θ*) = 0, the total effect is the comparative static

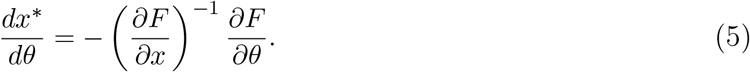

The inverse Jacobian couples all variables in a feedback component. The signs of its entries may determine a response direction, or unavailable derivative magnitudes may leave it unresolved.^31,32,40^ The PhysioMap solver returns ? in the latter case.

The relation types constrain the SCM differently. The PhysioMap solver projects the first-order derivative constraints for causal influence, production, and quantitative identity into a signed dependency graph *G*_∂_ = (*V, E*_∂_), whose signed adjacency pattern defines one signed Jacobian. Constitution does not enter the causal Jacobian; the PhysioMap solver represents its derivative signs in the constitutive dependency graph *G*_*κ*_ = (*V, C*) on the same vertices, with (*p*_*j*_, *w*) ∈ *C* labeled *σ*_*j*_ for each constituent in Table 1; adipose-tissue mass and lean-body mass each constitute total body weight. The constitutive dependency graph is acyclic because scales are strictly hierarchical, so no trait constitutes one of its own constituents, and determination is transferred along it in topological order by

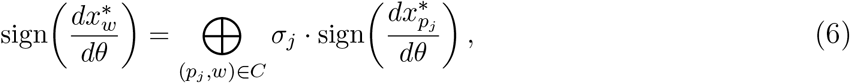

Where and are the sign product and parallel combination defined below, and a constituted variable that also carries a causal sign combines the two the same way. Because *C* has no edge (*w, p*_*j*_), a change in a whole is never propagated back to its parts. The signed dependency graph can contain cycles, so computing downstream effects must account for feedback.

After intervention, we remove the incoming dependencies of the target from *G*_∂_, yielding 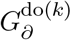, and decompose this graph into strongly connected components and process their acyclic condensation graph in topological order. A singleton uses sign algebra over {+, `-`, `0`, `?`} with `+ · +` = `- · -` = +, `+ · -` = `-`, and 0 annihilating products. An unknown factor absorbs a non-zero factor. The sign of a chain is the product of its edge signs. For parallel contributions, 0 is neutral, agreeing non-zero signs retain that sign, and opposing or unknown contributions give `?`. For a feedback component *S*, let *J*_*S*_ be its Jacobian sign pattern and *b*_*S*_ its signed forcing. The implicit function theorem and Cramer’s rule give

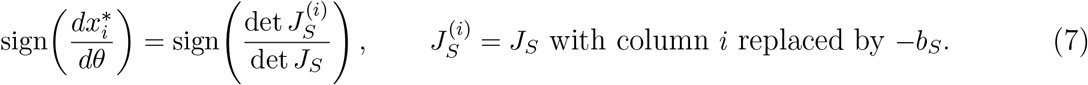

This links the derivative-sign abstraction to the quantitative comparative static without selecting a path.^41–43^ Suppl. Sec. S12.2 derives Eq. (7) from the component equilibrium equations and proves selective soundness. The exact component algorithm’s configurable size limit defaults to 16 traits. This bounds each exact determinant expansion by at most 2^16^ distinct subproblems and was fixed before evaluation as an engineering bound, not as a semantic or fitted threshold. The PhysioMap solver applies a conservative fixed-point algorithm to larger components; both algorithms abstain when a sign remains unresolved.

Figure 1 shows the representation and intervention steps in one CKG neighbourhood. Reducing plasma ACE activity removes its incoming causal influences and perturbs coupled volume and vascular-resistance pathways. Renal and baroreflex mechanisms close a whole-body feedback loop, while an intracellular arm connects receptor signals to smooth-muscle tone. The same neighbourhood contains a quantitative identity and a modulation relation, represented separately from causal influence.

**Fig. 1.**
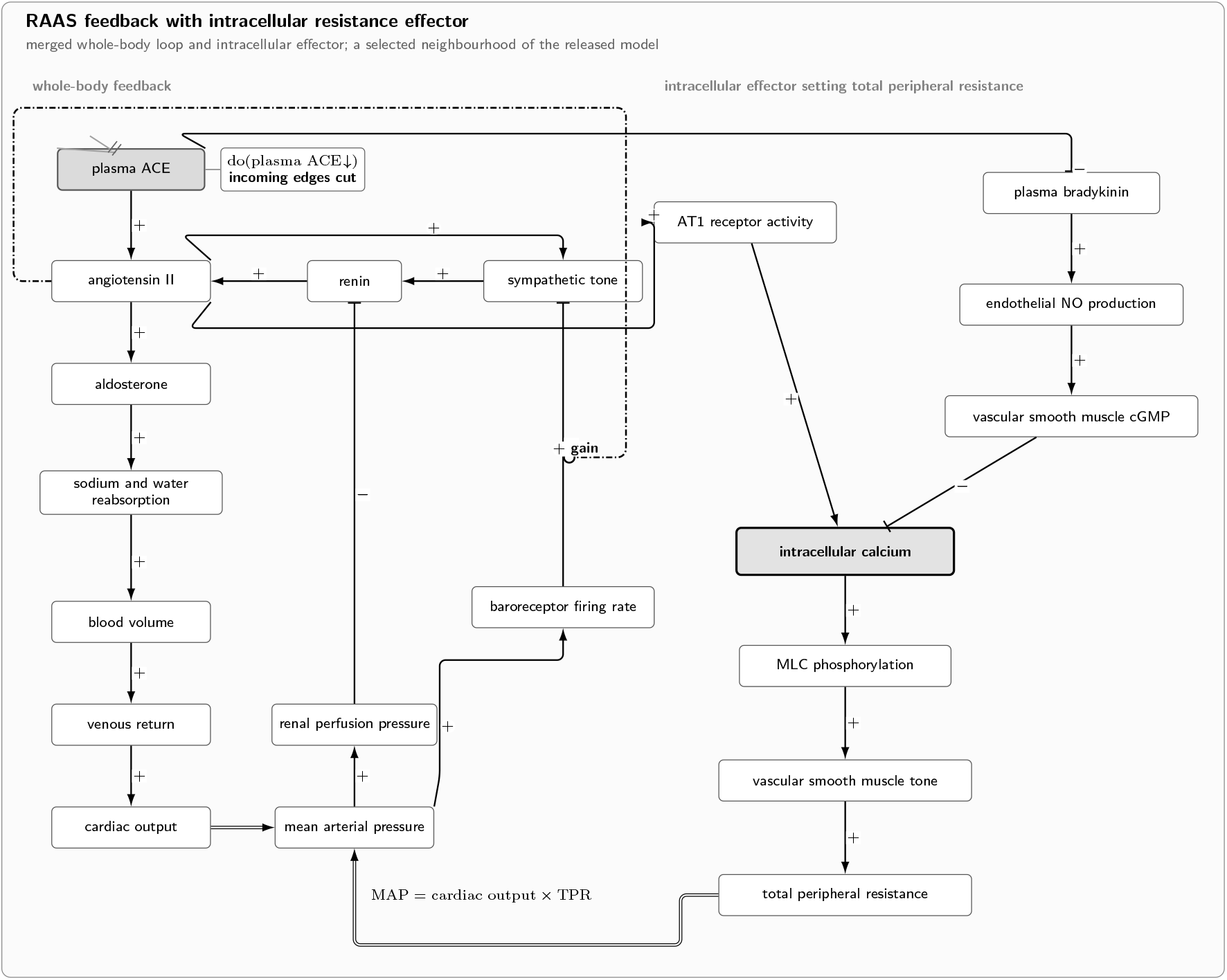
A selected PhysioMap neighbourhood merges whole-body renin–angiotensin–aldosterone system (RAAS) feedback with an intracellular resistance effector. Pointed arrows are positive causal influences; bars are negative influences. These glyphs show the corresponding SCM derivative signs. The double line denotes the quantitative identity mean arterial pressure (MAP) = cardiac output total × peripheral resistance (TPR), and the dash-dot line denotes modulation of the inhibitory baroreceptor-to-sympathetic influence. The intervention on plasma angiotensin-converting enzyme removes incoming causal influences. AT1, angiotensin II type 1 receptor; NO, nitric oxide; cGMP, cyclic guanosine monophosphate; MLC, myosin light chain.

### 2.5. Construction and content validation

We assembled PhysioMap by direct expert curation, extraction from quantitative models and curated databases, and checked language-model proposals from textbooks. The cardiovascular core used 17 Guyton BioModels modules and CellML integrators. Available equations supplied derivatives, while directional sources supplied constraints without complete functions. SIGNOR, AOP-Wiki, and direct literature curation provided additional content. Suppl. Sec. S15.1 lists the sources and acquisition metadata.

Following the entity–quality convention,^25–27^ each candidate trait received an entity identifier, PATO determinable, optional context, stable label, and scale. We resolved identifiers against pinned releases, classified measurement kind as extensive, ratio, rate, or intensive, checked that bearer and quality categories matched, and reconciled duplicates. Only composable measurement kinds admit part-to-whole rules.^44,45^ Unresolved traits remained explicitly primitive.

We assigned each relation to one of the five semantic types. Release checks covered schema, references, identifiers, bearers, constitution, relation typing, and regression against a frozen behavioral baseline. They assess formal consistency, not the scientific correctness of every axiom.

For cross-type content validation, P.N.S. reviewed a fixed-seed stratified sample of 83 v1.1.1 relations covering all five types. We interpreted `TRUE` as accepted, `FALSE?` as flagged for further review or investigation, and `FALSE` as rejected. Unequal sampling rates preclude a map-wide accuracy estimate. Suppl. Sec. S15.3 gives the protocol, sampling strata, and archive.

### 2.6. Evidence and provenance model

Evidence and provenance governed admission and documented construction; they are not PhysioMap content. Type, arguments, and context determine a relation’s semantics, and its sign constrains a derivative. Neither the reasoner nor the PhysioMap solver uses evidence class or provenance.

For causal-influence candidates, the evidence model admits six classes. Perturbation, pharmacological intervention, genetic loss or gain of function, and Mendelian randomization form the interventional group because they use a manipulation or valid instrument. Mechanistic-model evidence derives a sign from a curated model, while curated-mechanistic evidence records a curated functional or mechanistic account; these form the mechanistic group. Association, coexpression, or binding alone is insufficient. Other relation types use type-specific evidence, and historical support not yet mapped to a controlled class remains unclassified. Each stable relation identifier links to support and a projection trace.

### 2.7. Evaluation of prediction and abduction

We evaluated forward prediction against the HPO ontology and gene-to-phenotype annotation corpus (HPOA) from release 2026-02-16. For each gene entering the evaluation, we specified one or more primary lesion variables from gene-product identity and the disease mechanism. Suppl. Sec. S15.4 gives the mapping rule and links the complete versioned specification.

HPO phenotype classes follow an entity–quality (EQ) pattern, in which the entity *E* identifies the affected entity or process and the quality *Q* is drawn from PATO.^25,46^ This pattern lets us identify a directional HPO class when *Q* states an increase or decrease relative to normal. For example, hypoglycemia pairs blood glucose with decreased concentration (PATO:0001163). We encode the PATO distinction between increased quality (PATO:0002300) and decreased quality (PATO:0002301) as *s* = + and *s* = `-`, respectively.

We propagated HPOA annotations only upward through the HPO subclass hierarchy. Let *A*_*g*_ be the HPO classes directly annotated to gene *g*, let `HPO` denote the HPO ontology, and let *M* (*a*) = (*X*_*i*_, *s*) map a directional HPO class *a* to PhysioMap variable *X*_*i*_ and direction *s*. The reference directions for *X*_*i*_ were

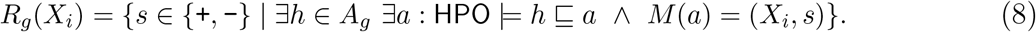

Therefore, a specific annotation *h* contributed the mapping of an ancestor *a*, never that of a descendant. We retained a gene–variable direction only when *R*_*g*_(*X*_*i*_) was a singleton. We blocked propagation for two deoxycortisol terms whose asserted HPO ancestry would otherwise map them to cortisol, although neither term denotes cortisol (Suppl. Sec. S15.4). We did not score the intervened variable.

Before scoring, we reviewed reference directions that disagreed with a PhysioMap solver prediction against the cited literature. Any resulting exclusion was applied uniformly to every method and reported as post hoc. Because HPOA and PhysioMap may draw on the same literature, we also repeated the analysis after excluding genes named in phenotype-directed curation fragments. Suppl. Sec. S15.4 gives the sources, adjudication rules, and overlap specification.

For each lesion, we applied the PhysioMap solver once and compared every downstream sign with the fixed reference pairs. Let *C, W*, and *A* denote correct, wrong, and abstaining pairs; missing and `?` predictions both count in *A*. We report precision = *C/*(*C* + *W*) and coverage = (*C* + *W*)*/*(*C* + *W* + *A*).

We compared two alternative inference rules on the same axioms and interventions. These rules differ only in how they combine directions, so the comparison isolates the contribution of feedback-aware inference rather than comparing independent phenotype predictors. The shortest-path rule receives a lesion and an endpoint, selects one shortest directed path, and multiplies its edge signs. Signed diffusion receives a lesion and assigns a score to every downstream trait. Let *S* be the signed, column-normalized adjacency matrix, so a node with out-degree *d* contributes +1*/d* or −1*/d* to each successor, and let *e* encode the signed intervention. We solved

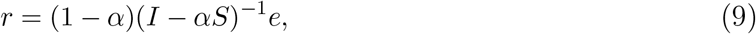

taking sign(*r*_*i*_) as the prediction and |*r*_*i*_| as its confidence. We set *α* = 0.85 and swept every distinct confidence observed on the evaluation pairs to produce the precision–coverage curve.

To test whether abstention identified cases that were difficult for the shortest-path rule without treating phenotype pairs from the same gene as independent, we used an exact within-gene conditional permutation test. Among pairs for which the shortest-path rule returned a direction, we held fixed each gene’s numbers of path errors and PhysioMap abstentions and permuted the abstention labels within that gene. The test statistic was the total number of path errors assigned to abstained pairs, and we report its exact one-sided upper-tail probability. Suppl. Sec. S15.4 gives the formal null distribution.

To test whether relation typing changed either operation, we repeated both evaluations while cumulatively adding production, quantitative identity, and constitution to causal influence. We compared forward predictions and best-tied inverse ranks with the complete first-order relation set. Modulation was excluded because these tasks score endpoint directions rather than mixed derivatives.

For abduction, we precomputed determinate phenotype signs for a fixed closed pool of single-lesion hypotheses and compared them with each observed directional profile. Agreement added one, contradiction subtracted one, and abstention scored zero. We ranked by net agreement, then fewer contradictions, then more agreements. Rank is one plus the number of strictly better candidates, so top-*k*, median rank, and mean reciprocal rank use the best position within a tie; unique top-1 requires no tie. We excluded the intervened trait and retained the true lesion in the pool.

Experiments used PhysioMap v1.1.1 and its locked dependencies. Suppl. Sec. S15.4 gives input checksums and file names; Suppl. Sec. S15.5 gives the reproduction commands.

## 3. Results

### 3.1. Construction, projection, and expert validation of PhysioMap

We built PhysioMap v1.1.1 as an OWL knowledge base containing 1,699 contextualized physiological traits and 2,270 causal-influence, 85 production, 4 constitution, 9 quantitative-identity, and 19 modulation axioms. Each axiom carries the evidence and provenance metadata described in Section 2.6; those annotations do not determine its relation type or application semantics.

From the entailed axioms, the versioned projection produced a typed causal knowledge graph with 1,699 trait nodes and 2,387 projected relation instances, represented as typed edges or hyperedges, across five relation types. This graph specifies a class of quantitative SCMs rather than a complete parameterized model or population distribution; many functions are constrained only by derivative signs. Its largest strongly connected component contains 213 traits. The PhysioMap solver reads the first-order derivative-sign abstraction and returns ? when it does not resolve an intervention response.

We validated the released content with the cross-type expert review. The expert accepted 69 of 83 sampled relations, flagged 12 for further investigation, and rejected 2 (Suppl. Table S1). Flagged relations were not counted as rejected, and both rejections were causal. The review therefore supports most sampled relations while identifying content that requires correction or further investigation.

### 3.2 Forward prediction recovers directional endophenotypes

The HPO mapping produced downstream lesion–phenotype directions for 167 of 183 matched genes; the other 16 genes had no mapped direction beyond the intervened variable. Before exclusion, the PhysioMap solver made 175 determinate predictions: 171 agreed with the HPO-derived reference and four disagreed, giving 97.7% agreement. Post hoc discrepancy review found that three of the four directions conflicted with the cited primary literature and one varied among studies. We excluded these four directions uniformly for every method, leaving an adjudicated reference of 866 pairs. Suppl. Sec. S15.4 identifies the pairs and gives the sources and rationales.

Against the adjudicated reference from HPOA release 2026-02-16, the PhysioMap solver returned 171 determinate directions, of which 171 agreed and 0 disagreed, and abstained on 695 pairs. This gives 100.0% precision at 19.7% coverage on the adjudicated reference set (Figure 2b). After we excluded the 15 genes with the most direct curation overlap, all 151 determinate directions remained concordant. These results support the joint behavior of the encoded dependencies and the PhysioMap solver, but they do not validate either component independently.

**Fig. 2.**
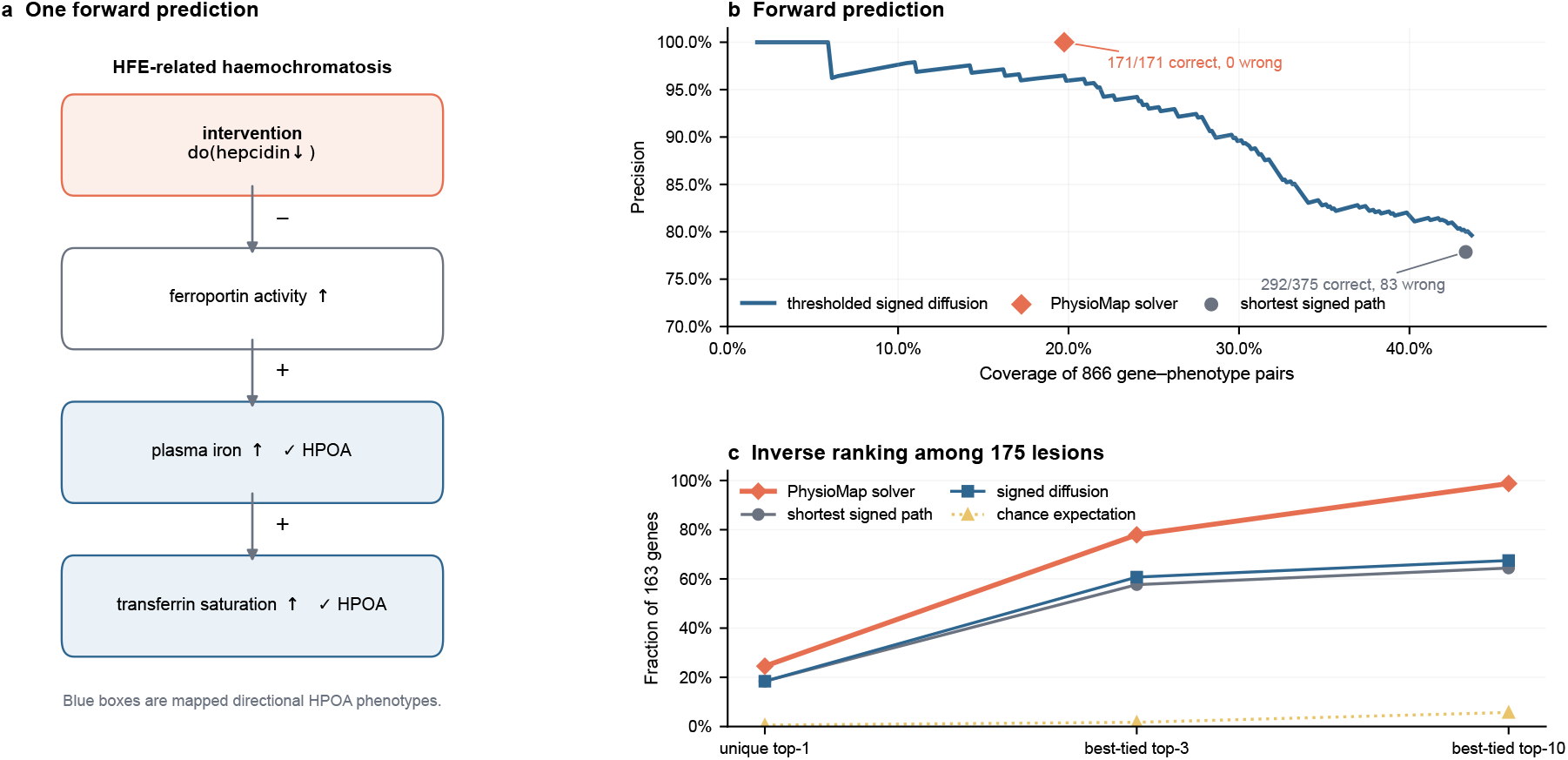
Evaluation of prediction and abduction. (a) Relations supporting an HFE-associated hereditary haemochromatosis prediction; only downstream phenotypes enter scoring. (b) Precision versus coverage for thresholded signed diffusion, with operating points and exact counts for the PhysioMap solver and shortest signed path. (c) Abductive ranking within 175 known lesions for the three inference rules and chance expectation. Top-3 and top-10 use the best rank within ties, and a unique top-1 requires no tied candidate; mean reciprocal rank for the PhysioMap solver is 0.707.

The shortest signed path returned 375 directions, of which 292 agreed and 83 disagreed with the reference, giving 77.9% precision. At *α* = 0.85, signed diffusion retained errors both without a confidence threshold and when thresholded to the same number of predictions as the PhysioMap solver (Figure 2b).

On all 171 pairs where the PhysioMap solver returned a direction, the path rule returned the same direction and therefore had a 0.0% error rate. The path rule additionally returned a direction for 204 pairs on which the PhysioMap solver abstained; only 121 (59.3%) agreed with HPOA, an error rate of 40.7%. Of the genes with a path prediction, 32 contained both shared-commitment and abstention pairs, and 15 contributed non-degenerate within-gene permutations. The exact conditional test rejected independence between abstention and path correctness (*P* = 2.05 *×* 10^−7^), showing that abstentions concentrated cases that were difficult for path-based inference after accounting for gene. This result does not show that every individual abstention was necessary or that the PhysioMap solver improves predictions when both methods commit.

The HFE trace shows how reduced hepcidin propagates through ferroportin to iron and transferrin-saturation traits (Figure 2a).

### 3.3. Abduction ranks candidate lesions from phenotype profiles

The inverse benchmark requires one primary lesion variable per gene. Four of the 167 genes with downstream reference directions, ARSA, CTNS, KYNU, and PKLR, each had two primary lesion variables and were therefore excluded, leaving 163 genes. For these genes, we ranked a closed pool of 175 single-lesion hypotheses that always included the true lesion. We held the pool, scoring rule, and tie handling fixed and changed only the inference rule that generated each candidate’s profile (Table 2).

**Table 2.** Inverse lesion ranking over 175 candidates for 163 genes. Scoring, tie handling, and metrics are identical in every row; only the rule producing a candidate’s predicted profile differs. Top-3 and top-10 use the best rank within ties, and a unique top-1 requires no tied candidate.

| inference rule | unique top-1 | top-3 | top-10 | MRR |
| --- | --- | --- | --- | --- |
| PhysioMap solver | 40 | 127 | 161 | 0.707 |
| shortest signed path | 30 | 94 | 105 | 0.535 |
| signed diffusion | 30 | 99 | 110 | 0.558 |
| chance | 1 | 3 | 9 | 0.033 |

All three inference rules exceeded the chance expectation. The PhysioMap solver placed the true lesion within the best-tied top three for 127 genes, compared with 94 for shortest path and 99 for diffusion, and had mean reciprocal rank 0.707 rather than 0.535 and 0.558. An abstention scores zero whereas an incorrect direction subtracts one, so withholding an unresolved direction can improve a candidate’s rank (Figure 2c).

The profiles narrowed the pool more reliably than they identified one lesion: the true lesion was uniquely first for only 40 of 163 genes. Because the closed pool always contains it, these results evaluate ranking among known lesions rather than open-set clinical diagnosis. The best position within a tie also makes top-*k* and reciprocal-rank results optimistic for every inference rule.

Across both applications, the complete first-order relation set changed four of 866 forward calls without a sign flip and 12 of 163 inverse ranks relative to causal influence alone, with no aggregate improvement (Suppl. Table S2). Numerical or population inference additionally requires functions, parameters, and exogenous distributions that PhysioMap v1.1.1 does not specify.

## 4. Discussion

PhysioMap provides a semantic layer between expert physiological maps and executable models. Ontology-grounded traits define what variables denote, while relation-specific SCM constraints define their dependencies. This permits intervention, abduction, and quantitative simulation without treating signs as the underlying model. The rare-disease application evaluates one use of this resource: the PhysioMap solver identified a set of unresolved cases on which the shortest-path baseline was substantially less reliable. This finding supports informative limits on qualitative prediction. The broader contribution is the semantic resource that makes this and other forms of causal inference possible.

The NIH Whole Person Initiative aims to map healthy organ-system functions, link them to common data elements, and construct an in silico prototype.^18,19^ PhysioMap could supply machine-interpretable trait identifiers and dependency semantics, while links to Physiome, CellML, and BioModels could attach reusable quantitative equations to the same traits;^12–14^ these integrations remain prospective.

Scaling requires semantic alignment, model composition, quantitative completion, and hybrid inference. Alignment must map measures and model variables to contextualized traits across anatomy, biological state, age, sex, time scale, and units. Composition must reconcile subsystem boundaries, conservation laws, incompatible time scales, and conflicting mechanisms. Quantitative completion must add functions, parameters, distributions, and uncertainty, with identifiability analysis. Solvers must combine sign constraints with numerical modules, decompose large feedback systems, and evaluate modulation and population effects. Versioned releases also require consistency checks, targeted expert review, model credibility assessment, and benchmarks.

The present evidence establishes a proof of concept, not clinical readiness, because expert sampling was unequal, the HPO-derived reference shares literature with PhysioMap, and abduction used a closed pool. Independent perturbation and drug-response data, prospective cohorts, open-set diagnosis, and comparisons with quantitative simulations across age, sex, and disease contexts are needed to test whether interoperability yields reproducible whole-person predictions.

