## Supplementary Information for "PhysioMap: an ontology-grounded causal knowledge graph of human physiology"

This supplement extends the core semantics stated in the main paper with complete OWL definitions, structural causal model and population semantics, the derivative-sign abstraction, the selective-soundness argument for the PhysioMap solver, and a generated characterization of release v1.1.1. PhysioMap's authored form is an OWL knowledge base. OWL classification identifies physiological trait classes and entailed relational patterns. A fixed projection assigns selected patterns to quantitative random variables, causal mechanisms, process-output production relations, exact functions, constitutive maps, and second-order modulation constraints. These constructs specify a class of admissible structural causal and population models; the current release does not instantiate all of their functions or distributions. Signs of first and second derivatives form a derived abstraction of this model class. The reported prediction and abduction experiments use the PhysioMap solver, which operates only on the first-order derivative-sign abstraction defined below. Evidence and provenance metadata used to construct and audit this content are defined separately and do not enter these semantics.

#### S1 OWL knowledge base and trait classes

Let $\mathcal{K}$ be an OWL knowledge base over a signature containing classes for entities, processes, qualities, traits, and model relations. It uses biomedical classes from resources including Uberon, CL, GO, ChEBI, PR, and PATO [3, 6, 8, 14, 15, 21], while defining its own continuant/process distinction and relations where required. The theory does not depend on a particular upper ontology or relation ontology.

An OWL interpretation is

$$\mathcal{I} = (\Delta^{\mathcal{I}}, \cdot^{\mathcal{I}}),$$

with the ordinary first-order semantics of OWL class and property expressions [13]. We write $\mathcal{I} \models \mathcal{K}$ when $\mathcal{I}$ satisfies the knowledge base and $\mathcal{K} \models \varphi$ when every such interpretation satisfies $\varphi$.

##### S1.1 Qualities and complete traits

PATO supplies attribute and value classes. A PATO quality, including an entity-specific quality, need not identify a unique physiological variable. PhysioMap therefore uses two levels of definition.

First, it post-composes a quality for entity $E_i$ and PATO quality $Q_i$:

$$\mathsf{EntitySpecificQuality}_i \sqsubseteq \mathsf{Quality} \sqcap Q_i \sqcap \exists\, \mathsf{concerns}.E_i. \tag{S1}$$

Second, a complete trait states that the organism has a part which is the characterized entity or process, sits in the recorded context, and bears that quality:

$$T_i \sqsubseteq \mathsf{Trait} \sqcap \exists\, \mathsf{hasPart}.\big(E_i \sqcap \exists\, \kappa_i.C_i \sqcap \exists\, \mathsf{hasQuality}.\mathsf{EntitySpecificQuality}_i\big), \tag{S2}$$

with $\kappa_i = \mathsf{contextPartOf}$ for a continuant entity and $\kappa_i = \mathsf{occursIn}$ for a process; the context conjunct is dropped when no context $C_i$ is recorded. This is the EQ principle used throughout PhysioMap [6, 7], stated so that the bearer of the quality is a part of the organism rather than an attribute of the trait class itself. hasContinuantPart and hasOccurrentPart are the subproperties of hasPart actually asserted, matching the continuant/process split of the trait. Additional restrictions may specify cell type, participants, temporal context, measurement kind, or scale.

#### S1.2 Mechanism relations and the collection pattern

A causal, production, or modulation axiom is a type-level relation claim. This is its formal content, irrespective of the evidence and provenance metadata used to admit and document it. The subclass form $T_t \sqsubseteq \exists\,\mathsf{causedBy}.T_s$ states something stronger and generally false for the intended type-level claim: that every bearer of the target trait is caused by an instance of the source. One organism in which the source arm is absent or pharmacologically blocked, while the target trait persists, refutes it. The faithful dispositional reading $T_t \sqsubseteq \exists\,\mathsf{hasDisposition}.(\forall\,\mathsf{realizedBy}.\exists\,\mathsf{hasParticipant}.T_s)$ nests a universal inside an existential restriction, leaves OWL 2 EL, and is not classifiable at this scale [10].

PhysioMap therefore uses the collection pattern developed for biological function [10]. Every trait $T$ has an associated collection class $C_T$, with

$$T \sqsubseteq \exists\,\mathsf{memberOf}.C_T, \qquad C_T \sqsubseteq \exists\,\mathsf{hasMember}.T, \tag{S3}$$

and a mechanism axiom from $T_s$ to $T_t$ asserts

$$C_{T_t} \sqsubseteq \exists\,\mathsf{hasMember}.(\exists\,\mathsf{causedBy}.T_s), \tag{S4}$$

that is, some member of the target's collection is actually caused by an instance of the source, which supplies the existential witness required by the type-level claim. Production and modulation take the same shape with producedBy and modulates. Constitution does not, because a whole's extensive trait is constituted by its parts' traits in every instance; it keeps $T_{\text{macro}} \sqsubseteq \exists\,\mathsf{constitutedBy}.T_{\text{micro}}$.

Two consequences are worth stating. First, all collection-pattern axioms above are existential, so the artifact ELK classifies is unchanged in expressivity and the released entailments are unchanged in content. Second, inheritance becomes asymmetric, and correctly so: a member caused by a $T_s$ instance is caused by an instance of every superclass of $T_s$, so the source generalizes, but the witness member need not fall under a more specific target trait, so the target does not specialize. The subclass form licensed that second inference and should not have.

Collections are not map variables and never enter the SCM; each is tied to its trait by a collectionFor annotation carrying the node identifier, so no semantics depends on the lexical form of a PhysioMap IRI.

The released OWL artifacts separate what PhysioMap represents from what it reasons over. The EL artifact carries the axioms above together with the chain $\mathsf{occursIn} \circ \mathsf{partOf} \sqsubseteq \mathsf{occursIn}$, transitivity of partOf, and the part-inclusive grouping classes; ELK classifies it and answers every released query. The mirror chain $\mathsf{hasPart} \circ \mathsf{partOf} \sqsubseteq \mathsf{hasPart}$ is deliberately absent: it is unsound, since a part of my part's whole need not be my part, and with partOf as the inverse of hasPart it also breaks OWL 2 DL regularity; the part-inclusive closure runs through reflexive partOf helper classes instead. The DL artifact adds the axioms outside OWL 2 EL: the inverses partOf, qualityOf, siteOfProcess, and memberOf, the domains and ranges of localization, PATO's universal constraint that a process quality inheres only in processes, $\mathsf{ProcessQuality} \sqsubseteq \forall\,\mathsf{qualityOf}.\mathsf{Process}$, and the two commitments about collections that EL cannot state: homogeneity $C_T \sqsubseteq \forall\,\mathsf{hasMember}.T$ and the singleton $C_T \equiv \{c_T\}$. Each external entity is typed as a continuant or an occurrent. The released reasoning workflow never reads the DL artifact, so representational commitments cannot silently change a reported entailment.

A directional HPO value class and a causal change remain distinct. Membership in an increased-concentration phenotype is relative to a reference range. A response $\Delta X > 0$ is relative to the pre-intervention value and need not cross that range.

#### S2 One population space and trait-valued random variables

A population instantiation of PhysioMap is based on one probability space

$$(\Omega, \Sigma, P).$$

All nodes in one SCM instantiation use this same space. An element $\omega \in \Omega$ is a sampling unit containing sufficient subject, time, and physiological context to assign the traits in the instantiation. Using one space gives a joint distribution, rather than a collection of unrelated marginals.

Let

$$\mathcal{V} = \{T_1, \ldots, T_n\}$$

be the finite set of complete trait classes selected as model variables. Each $T_i$ is associated with an ordered measurable value space $(D_i, \mathcal{D}_i, \leq_i)$ and a measurable random variable

$$X_i : (\Omega, \Sigma) \longrightarrow (D_i, \mathcal{D}_i).$$

Together they define

$$X = (X_1, \ldots, X_n) : \Omega \longrightarrow \prod_i D_i$$

and the joint distribution $P_X = P \circ X^{-1}$. The marginal for a node is $P_i = P \circ X_i^{-1}$.

A population instantiation assigns one population jointly to all nodes, rather than a separate population to each node. Different populations give different instantiations of the same structural model, for example

$$(\mathcal{M}, P_{\text{UKB}}) \quad \text{and} \quad (\mathcal{M}, P_{\text{MIMIC}}),$$

or conditional distributions $P(\cdot \mid S_d = 1)$ under explicit selection variables.

##### S2.1 The realization and valuation maps

The OWL domain $\Delta^{\mathcal{I}}$ and the sampling population $\Omega$ are not identified. For every selected trait define

$$\rho_i : \Omega \longrightarrow T_i^{\mathcal{I}}$$

to select the trait instance realized by a sampling unit, and a measurable valuation

$$\nu_i : T_i^{\mathcal{I}} \longrightarrow D_i.$$

Compatibility requires

$$X_i = \nu_i \circ \rho_i. \tag{S5}$$

This is the precise meaning of the statement that a trait class becomes a random variable. The OWL class supplies the type and logical constraints; $X_i$ supplies the population value.

For a determinate class $H \sqsubseteq T_i$, define

$$A_{i,H} = \{\omega \in \Omega : \rho_i(\omega) \in H^{\mathcal{I}}\}.$$

These events are required to belong to $\Sigma$. Standard OWL consequences then constrain the population measure:

$$\mathcal{K} \models H \sqsubseteq H' \implies A_{i,H} \subseteq A_{i,H'} \implies P(A_{i,H}) \leq P(A_{i,H'}), \tag{S6}$$

$$\mathcal{K} \models H \equiv H' \implies A_{i,H} = A_{i,H'}, \tag{S7}$$

$$\mathcal{K} \models H \sqcap H' \sqsubseteq \bot \implies A_{i,H} \cap A_{i,H'} = \varnothing. \tag{S8}$$

The binary causal-knowledge-graph semantics of Toonsi et al. [22] is recovered by the indicator random variable $\mathbf{1}_{A_{i,H}}$. PhysioMap generalizes this construction to ordered and continuous traits.

For an intervention regime $a$, the same construction uses $X_i^a$ and a regime-indexed realization $\rho_i^a$. It is sufficient either to use regime-indexed trait instances in one OWL domain or a family of interpretations $\mathcal{I}^a \models \mathcal{K}$. In either case, the three entailment constraints above hold in every regime.

#### S3 Entailment-to-model projection

OWL does not express structural equations or the do-operator. PhysioMap therefore supplies a finite, versioned library $\Pi$ of projection rules. A rule $p \in \Pi$ has an OWL entailment premise and a quantitative structural causal model conclusion:

$$\mathcal{K} \models \varphi_p(T_{i_1}, \ldots, T_{i_k}) \quad \Longrightarrow \quad \psi_p(X_{i_1}, \ldots, X_{i_k}). \tag{S9}$$

The rules are defined over classified entailments [9, 18]. Consequently, entailed subclasses, equivalences, parthood relations, and other relations may change which relational patterns match. In release v1.1.1, typed relations are encoded losslessly in OWL and the release validation uses ELK to verify their registered relational witnesses.

We call the materialized typed relation structure the causal knowledge graph (CKG). It is distinct from both the OWL knowledge base and the signed dependency graph derived for first-order computation. Projection is intended to produce the CKG's quantitative content specification

$$\pi_{\Pi}^{\text{content}}(\mathcal{K}) = (V, E, \mathcal{P}, \mathcal{C}, \mathcal{Q}, \mathcal{H}, \mathcal{F}, \Phi), \tag{S10}$$

where:

- $V$ contains the named trait classes, including explicit incomplete primitives;
- $E$ contains the contextualized causal influences whose source–target pairs form the ordinary causal parent graph;
- $\mathcal{P}$ contains typed process-output or consumption relations;
- $\mathcal{C}$ contains constitutive constraints and $\mathcal{Q}$ contains quantitative expressions;
- $\mathcal{H} \subseteq V \times E$ contains modulation relations that constrain second derivatives;
- $\mathcal{F}$ constrains the admissible quantitative causal mechanisms;
- $\Phi$ contains exact measurement and constitutive functions.

The release manifest materializes $V, E, \mathcal{P}, \mathcal{C}, \mathcal{Q}, \mathcal{H}$. $\mathcal{F}$ and $\Phi$ denote the functions admitted by those projected objects; they are not complete numerical mechanism libraries serialized in release v1.1.1. The materialized relation objects may carry first- or second-derivative sign constraints.

These constraints restrict $\mathcal{F}$ and $\Phi$, but they do not replace the quantitative functions or random variables in the semantics. Section S11 defines the derivative-sign abstraction derived from this model class.

The authoritative OWL file contains a lossless encoding of every projected content object. This includes the elements of $V$ and $E$ and the typed objects in $\mathcal{P}, \mathcal{C}, \mathcal{Q}, \mathcal{H}$. Every typed relation has a stable identifier. A projected object is admitted to the release only if it has its registered OWL witness.

A separate map $\tau_\Pi(\mathcal{K})$ links each stable content identifier to its evidence and provenance metadata. This registry records the entailed relational witness, supporting source axioms, projection pattern and version, reasoner, and source ontology versions, as well as relation-level evidence annotations where applicable. $\tau_\Pi(\mathcal{K})$ is not a component of $\pi_\Pi^{\text{content}}(\mathcal{K})$, does not constrain $\mathfrak{M}(\mathcal{K}, \Pi)$, and is not an input to inference.

The projected CKG is closed only relative to the declared PhysioMap model boundary. OWL remains open-world: the absence of an OWL assertion does not imply that no real cause exists. For an SCM, however, $\text{pa}(i)$ is the complete set of modeled direct parents of $X_i$; unmodeled influences are absorbed by exogenous or boundary variables.

### S4 Reasoning boundary and release artifacts

The public release contains `physiomap.owl`, the complete TBox-only primary knowledge base with signature-bound source modules; `physiomap-el.owl`, the enforced OWL 2 EL reasoning artifact; the versioned projection registry; `physiomap-scm.json`; and separate projection traces and migration reports. No population member or other named ABox individual is copied into the primary knowledge base. Source modules are extracted by OWLAPI bottom-locality extraction from checksum-pinned ontology releases.

In the fixed Functional Syntax serialization for release v1.1.1, `physiomap.owl` contains 126,344 top-level axioms. Of these, 36,788 are logical axioms; the remainder are declarations and annotations, including imported labels and embedded projection entries. These counts were obtained by grouping the checksum-pinned file by top-level OWL constructor. Checksums for the release artifacts are listed in the machine-readable release checksum manifest. The total excludes the `Prefix` declarations and `Ontology` wrapper; the logical count also excludes `Declaration` and `AnnotationAssertion` axioms.

OWLAPI first checks that `physiomap-el.owl` is in the OWL 2 EL profile. ELK then provides complete reasoning for that enforced EL artifact, including classification, consistency, trait satisfiability, and registered projection entailments. Non-EL validation is permitted only for explicitly registered locality modules with declared signatures, axiom limits, timeouts, memory bounds, and contradiction fixtures; HermiT is complete only for each such bounded OWL 2 DL module. We do not claim unrestricted full-knowledge-base OWL 2 DL consistency. An empty non-EL registry means that the current PhysioMap projection itself requires no non-EL check, not that arbitrary non-EL axioms are silently ignored.

Candidate substitutions come from asserted restrictions and their inferred class closures, so validation does not require all-pairs HermiT reasoning. Every release also compares the optimized ELK projection with the expected typed SCM constructs, verifies equivalence to the migration-source representation, realizes all declared first- and mixed-derivative signs numerically, and compares outputs from the PhysioMap solver, SCC decomposition, HPO evaluation, benchmarks, public interface, and website with versioned references.

In release v1.1.1, the OWL knowledge base, projection registry, and generated SCM are the authoritative release layers. Opposing insulin effects on VLDL secretion are retained as separate fed-state lipogenic and direct insulin-signaling contexts.

#### S5 Evidence and provenance model for construction

Evidence and provenance govern the admission and audit of candidate content. They are metadata about an axiom, not additional PhysioMap relation types. In particular, an evidence label cannot change a production relation into a causal influence, change the sign of an influence, or alter its intervention semantics. The annotations are ignored by OWL logical entailment and are not read by the PhysioMap solver.

For causal-influence candidates, PhysioMap recognizes six controlled evidence classes. Perturbation, pharmacological intervention, genetic loss or gain of function, and Mendelian randomization comprise the interventional group: each supports an effect through a manipulation or valid instrument. Mechanistic-model evidence derives a direction from the equations or derivatives of a curated quantitative model. Curated-mechanistic evidence records a causal claim supported by a curated functional or mechanistic account. These two classes comprise the mechanistic group. Therefore, *interventional* and *mechanistic* in release summaries classify the support for a causal axiom, not its content. Binding, coexpression, and observational association alone are insufficient to admit a causal influence.

Production uses a separate evidence vocabulary because its content is a process–output relation. Modulation requires support for the asserted interaction between influences. Constitution and quantitative identity are admitted through explicit mereological or mathematical patterns. Historical causal relations whose support has not yet been mapped to a controlled class remain explicitly unclassified in the evidence model rather than being bulk reclassified. Each included relation is linked through $\tau_\Pi(\mathcal{K})$ to its supporting source or evidence and its projection trace.

#### S6 Structural causal model semantics

The intended semantics of the projection is a class $\mathfrak{M}(\mathcal{K}, \Pi)$ of quantitative structural dynamical causal models. This model class, rather than its derivative-sign abstraction, supplies the semantics of the projected variables and relations. A member contains exogenous random variables $U = (U_1, \dots, U_n)$ and twice continuously differentiable mechanisms

$$\dot{x}_i = F_i(x_i, x_{\mathrm{pa}(i)}, U_i), \qquad i = 1, \dots, n. \tag{S11}$$

The baseline population specification assumes a product distribution for the $U_i$, so it represents no latent common causes. This is a modeling assumption, not a property inferred from the OWL knowledge base; an extension may use dependent exogenous variables and bidirected edges.

An ordinary projected edge $j \to i$ means that $F_i$ functionally depends on $x_j$. Its direct quantitative effect at state $x$ and context $u$ is

$$g_{ji}(x, u) = \frac{\partial F_i}{\partial x_j}(x, u). \tag{S12}$$

Where the function is known, $\mathcal{F}$ may specify it. Where only an order constraint is known, $\mathcal{F}$ contains a class of functions, for example

$$\frac{\partial F_i}{\partial x_j} > 0 \quad \text{throughout an operating neighborhood.}$$

Therefore a relation sign is derived from, and constrains, a quantitative derivative. It is not the value of a physiological variable and does not replace the SCM semantics:

$$\sigma_{ji} = \mathrm{sign}(g_{ji}).$$

For every allowed regime and exogenous context, assume a locally unique equilibrium $x^{*,a}(u)$ on the physiological operating branch whose Jacobian is Hurwitz. Under the equilibration and intervention conditions of dynamical causal models, the equilibrium equations form a cyclic SCM [2, 12, 19]. The population random variables are then

$$X_i^a(\omega) = x_i^{*,a}(U(\omega)). \tag{S13}$$

##### S6.1 Direct-effect identity

Fix distinct variables $s$ and $t$, an exogenous context $u$, and a reference equilibrium $x^*$. Under $\mathrm{do}(x_s = \theta,\ x_j = x_j^* \text{ for } j \notin \{s,t\})$, only $x_t$ is allowed to re-equilibrate. Write

$$G(z,\theta) = F_t\big(z, \theta, x^*_{-(s,t)}, u_t\big).$$

At the reference point, $G(x_t^*, x_s^*) = 0$. Local self-regulation gives

$$\left.\frac{\partial G}{\partial z}\right|_{(x_t^*, x_s^*)} = \left.\frac{\partial F_t}{\partial x_t}\right|_{(x^*,u)} < 0,$$

so this derivative is non-zero. The implicit function theorem therefore gives a locally unique differentiable function $h$ such that $h(x_s^*) = x_t^*$ and $G(h(\theta), \theta) = 0$. Differentiating the latter identity at $\theta = x_s^*$ gives

$$0 = \frac{\partial F_t}{\partial x_t} h'(x_s^*) + \frac{\partial F_t}{\partial x_s}, \qquad h'(x_s^*) = -\frac{\partial F_t / \partial x_s}{\partial F_t / \partial x_t},$$

where both partial derivatives are evaluated at $(x^*, u)$. By the definition of the direct intervention effect, $D_{st} = h'(x_s^*)$. Since $-1/(\partial F_t/\partial x_t) > 0$, multiplication by this factor does not change a sign. Hence

$$\mathrm{sign}(D_{st}) = \mathrm{sign}\left(\frac{\partial F_t}{\partial x_s}\right).$$

This proves the direct-effect identity used in the main paper.

#### S7 Measurement and constitutive projection rules

OWL parthood or PATO typing alone does not determine a distribution. They affect the probabilistic model when a complete entailed pattern activates a bridge rule with an explicit measurable function. Exact rules hold $P$-almost surely in every regime in which their premises hold. The distribution of the target is the pushforward of the joint source distribution under that function.

PhysioMap classifies PATO attributes with a measurement-behavior overlay. The initial classes are extensive, ratio, rate, product-defined, monotone-derived, and intensive/unconstrained. This is a classification for projection; it does not modify PATO.

#### S7.1 Production and consumption

A production relation is a typed process-output statement: a process produces, secretes, or releases an output, or consumes/removes it. It does not assert parthood and is not stored in $E$ as an authored causal influence. Its sign is a derivative abstraction of the quantitative constraint. For a production relation $(p, t, \sigma)$, an admissible model has

$$F_t = R_t + \epsilon_{pt} q_{pt}(x_p), \qquad q'_{pt} > 0, \qquad \epsilon_{pt} \in \{-1, +1\}, \qquad \epsilon_{pt} = \sigma,$$

where $R_t$ is independent of $x_p$. The direct-effect identity then gives $\mathrm{sign}(D_{pt}) = \sigma$. For the PhysioMap solver, projection derives the signed dependency

$$(p, t, \sigma) \in \mathcal{P} \quad \longmapsto \quad p \xrightarrow{\sigma} t.$$

The canonical relation remains production; only its derived first-order constraint enters the PhysioMap solver.

#### S7.2 Extensive attributes

Suppose $\mathcal{K}$ entails that entities $E_1, \ldots, E_k$ are pairwise disjoint and jointly exhaustive parts of $E$, and that $T_1, \ldots, T_k, T$ carry the same extensive attribute. The projection rule is

$$X_T = \sum_{r=1}^{k} X_{T_r} \qquad P\text{-almost surely.} \tag{S14}$$

This applies to amounts, masses, volumes, counts, and other quantities with an empirical concatenation operation admitting an additive representation [11, 20]. It follows that

$$\frac{\partial X_T}{\partial X_{T_r}} = 1.$$

Neither parthood without completeness nor an intensive attribute activates Eq. (S14). Dependence among the parts is retained: the distribution of a sum is determined by their joint distribution, not their marginals alone.

#### S7.3 Ratio attributes and optional decomposition

A ratio trait has quantitative semantics on a positive domain

$$X_R = \frac{X_N}{X_D}, \qquad X_N > 0, \quad X_D > 0, \tag{S15}$$

where $N$ is a numerator quantity and $D$ a denominator quantity. Concentration, density, fractions, and normalized indices are examples [11, 17]. The defining derivatives are

$$\frac{\partial X_R}{\partial X_N} = \frac{1}{X_D} > 0, \qquad \frac{\partial X_R}{\partial X_D} = -\frac{X_N}{X_D^2} < 0. \tag{S16}$$

For positive values,

$$\frac{dX_R}{X_R} = \frac{dX_N}{X_N} - \frac{dX_D}{X_D}.$$

Hence simultaneous increases of numerator and denominator have no fixed sign without magnitudes.

The numerator and denominator may be explicit selected traits. In that case Eq. (S15) enters $\Phi$, and the corresponding dependencies and interventions are available to the SCM. If either is not selected, it remains an internal argument of the ratio's quantitative interpretation. The ratio node remains valid, but the hidden argument is not independently queryable or intervenable. This allows gradual decomposition without making it a requirement of the present resource.

For plasma glucose concentration, the numerator may be amount or mass of glucose in plasma and the denominator plasma volume or mass. The exact trait therefore also records the concentration kind and value unit; an amount concentration and a mass fraction are not the same variable even when values can be converted using additional information.

##### S7.4 Rates

A rate is a ratio of an accumulated extensive quantity $Q$ to duration $\tau$:

$$X_R = \frac{X_Q}{X_\tau}, \qquad X_\tau > 0. \tag{S17}$$

This covers flow, secretion, reaction, and event rates. When the numerator and duration are exposed and positive, Eq. (S16) applies. Aggregation of concurrent rates is justified only when their flows are composable over the same interval; a rate is not generically additive merely because PATO classifies it as a rate.

##### S7.5 Products and monotone transformations

A physical or definitional pattern may provide

$$X_T = \prod_{r=1}^{k} X_{T_r} \tag{S18}$$

on a stated domain. Examples include CO = SV × HR and an appropriate MAP = CO × TPR approximation. On a strictly positive domain, every partial derivative in Eq. (S18) is positive. These equations require an explicit physical-law or definition pattern and do not follow from a PATO attribute alone.

More generally, if an entailed pattern supplies $X_T = g(X_S)$, then its distribution is $P_{X_S} \circ g^{-1}$. The derivative sign follows from $g'$ on the operating domain. This covers increasing and decreasing transformations and unit conversions.

##### S7.6 Intensive attributes

Pressure, temperature, pH, activity, resistance, potential, and other intensive attributes have no generic part-to-whole equation. Their whole values may be weighted averages, nonlinear functions, equilibria, or quantities determined by mechanisms outside mereology. Projection therefore adds no distributional or causal constraint from parthood alone. A specific function or causal pattern is required.

#### S8 Constitution and causal abstraction

An exact constitutive equation in $\Phi$ is a measurable map

$$X_M^a = \phi_M(X_{m_1}^a, \ldots, X_{m_k}^a) \qquad P\text{-almost surely} \tag{S19}$$

for every allowed aligned regime $a$. It is non-causal: it relates two descriptions of the same system rather than two independently changeable mechanisms. Equation (S14) and an exposed Eq. (S15) are exact examples.

For intervention lifting between grains, a map from allowed micro interventions to macro interventions must make abstraction commute with intervention [1, 19]. If $\alpha = \phi_M$ is the state abstraction and $\omega$ maps interventions, the requirement is

$$\alpha_{\#} P^{a}_{\mathcal{M}_L} = P^{\omega(a)}_{\mathcal{M}_H}, \tag{S20}$$

where $\alpha_{\#}$ denotes pushforward. Macro-to-micro lifting is not generally defined because many micro configurations can realize the same macro value. A purported cross-scale relation without a complete $\phi_M$ is a structural or causal hypothesis, not exact constitution.

Exact constitutive equations do not enter the causal Jacobian. When their source traits are explicit, their target variables are deterministic nodes for separation purposes. If an allegedly constituted target also has non-definitional parents, the exact identity is incomplete and the node is conservatively treated as ordinary causal structure.

For observational separation, auxiliary functional-dependency edges connect the arguments of an exact function to its target. These dependencies state that the target value is fixed by its arguments; they are not authored causal claims or independently intervenable mechanisms. Their derivative signs form a separate, acyclic constitutive dependency graph. The PhysioMap solver uses this graph to transfer a determined change from explicit constituents to a constituted variable between causal components; constitutive dependencies do not enter the causal Jacobian or become causal edges.

### S9 Multiplicative modulation

#### S9.1 OWL representation

A modulation statement is about a causal influence, not only about its target. OWL object properties are binary, so PhysioMap represents the underlying causal influence as a first-class, reified pattern. At the semantic level, a causal-influence object $c$ identifies source trait $T_j$ and target trait $T_i$. A modulation object $q$ identifies modulator trait $T_m$, the influence $c$, and a signed mixed-derivative direction $\eta$:

$$c = \text{Influence}(T_j, T_i), \qquad q = \text{Modulation}(T_m, c, \eta).$$

This n-ary content can be encoded with ordinary OWL classes and binary properties such as hasSourceTrait, hasTargetTrait, hasModulatorTrait, and modulatesInfluence. Classification may infer the endpoint types, equivalence of influence patterns, or membership in a positive or negative modulation class. The quantitative meaning is supplied by the following projection rule, not by changing the ordinary OWL semantics of those properties.

#### S9.2 Projection and mixed-derivative abstraction

If classification entails

$$\mathcal{K} \models \text{Modulation}(T_m, \text{Influence}(T_j, T_i), \eta),$$

projection adds the hyperedge

$$(m, (j, i)) \in \mathcal{H}. \tag{S21}$$

The ordinary causal influence is the derivative

$$g_{ji}(x,u) = \frac{\partial F_i}{\partial x_j}(x,u).$$

The modulation sign abstracts the sign of the change in this quantitative slope:

$$\eta_{m;ji} = \operatorname{sign}\left(\frac{\partial g_{ji}}{\partial x_m}\right) = \operatorname{sign}\left(\frac{\partial^2 F_i}{\partial x_m\, \partial x_j}\right). \tag{S22}$$

Therefore, $\eta = +$ means that increasing the modulator makes the signed slope more positive; it does not necessarily increase the magnitude of the base influence.

To recover the magnitude interpretation in an operating neighborhood with a sign-stable base effect, factor the slope as

$$g_{ji}(x,u) = \sigma_{ji} k_{ji}(x_m, z, u), \qquad \sigma_{ji} \in \{-1,+1\}, \quad k_{ji} > 0, \tag{S23}$$

where $z$ denotes other context. The sign of the change in magnitude is

$$\lambda_{m;ji} = \operatorname{sign}\left(\frac{\partial k_{ji}}{\partial x_m}\right) = \sigma_{ji}\eta_{m;ji}. \tag{S24}$$

Consequently, $\lambda = +$ strengthens the magnitude and $\lambda = -$ weakens it. For a negative base influence, $\eta = +$ gives $\lambda = -$: the slope moves toward zero and the inhibition weakens. This is the convention used by the projection registry and numerical mixed-derivative validator.

The hyperedge in Eq. (S21) is retained as a second-order object. If $F_i$ functionally depends on $x_m$, the ordinary SCM graph must also contain $m \to i$, even if the first-order derivative happens to vanish at a centered operating point. SCM parenthood is defined by functional dependence, while $\mathcal{H}$ records that the scientifically intended role of that dependence is modulation of $j \to i$.

A modulation may be fully quantitative, with a specified $k_{ji}$, or partially quantitative, with a class of positive differentiable functions satisfying Eqs. (S22)–(S24). A sign-changing influence cannot use one global factorization with $k_{ji} > 0$; it must be divided into sign-stable regimes or explicitly marked as capable of crossing zero.

##### S9.3 Population interpretation

At the population operating points, edge gain and modulation strength are themselves measurable random variables:

$$G_{ji}(\omega) = \left.\frac{\partial F_i}{\partial x_j}\right|_{X(\omega),U(\omega)}, \tag{S25}$$

$$H_{m;ji}(\omega) = \left.\frac{\partial^2 F_i}{\partial x_m\, \partial x_j}\right|_{X(\omega),U(\omega)}. \tag{S26}$$

An asserted sign constraint is required to hold almost surely in the declared regime. Individuals may nevertheless differ in the magnitudes of $G_{ji}$ and $H_{m;ji}$.

##### S9.4 Local structural interaction and total equilibrium synergy

The mixed derivative in Eq. (S22) is local to one structural mechanism. The total interaction after feedback is a second derivative of the equilibrium. Let

$$F(x^*(\theta),\theta) = 0, \qquad J = F_x.$$

For intervention parameters $\theta_a, \theta_b$, implicit differentiation gives

$$x_a = -J^{-1}F_a, \tag{S27}$$

$$x_{ab} = -J^{-1}\left(F_{ab} + F_{xa}x_b + F_{xb}x_a + F_{xx}[x_a, x_b]\right). \tag{S28}$$

The structural modulation terms are entries of $F_{xx}$. Through $J^{-1}$, feedback and alternative paths can amplify, cancel, or reverse their contribution to $x_{ab}$. Therefore the sign $\eta_{m;ji}$ is the intrinsic structural interaction sign; it is not automatically the sign of whole-system synergy. They agree only when the remaining terms and propagation preserve it. Any method that infers this total interaction must evaluate or soundly abstract Eq. (S28).

### S10 Interventions, distributions, and stochastic order

A perfect intervention $\mathrm{do}(x_k = \theta)$ replaces the $k$th mechanism with the assignment $x_k = \theta$, deleting its incoming functional dependencies and leaving all other mechanisms invariant [16]. For the reduced equilibrium system,

$$\frac{dx^*_{-k}}{d\theta} = -[J^{(-k)}]^{-1}b^{(k)}, \tag{S29}$$

where $J^{(-k)}$ is the reduced Jacobian and $b^{(k)}$ the derivative of the remaining equations with respect to the clamped value. Under regular convergence, this equilibrium sensitivity is the $t \to \infty$ limit of the dynamical intervention response.

The exogenous distribution and structural mechanisms induce an interventional random vector $X^a$ and distribution $P^a = P \circ (X^a)^{-1}$. A derivative sign and a population stochastic ordering are related, but not identical. If

$$X_i^{\mathrm{do}(x_k=\theta_2)}(\omega) \geq X_i^{\mathrm{do}(x_k=\theta_1)}(\omega) \quad \text{for almost every } \omega, \qquad \theta_2 > \theta_1,$$

then

$$P\Big(X_i^{\mathrm{do}(x_k=\theta_2)} > z\Big) \geq P\Big(X_i^{\mathrm{do}(x_k=\theta_1)} > z\Big) \quad \text{for every } z. \tag{S30}$$

This is first-order stochastic dominance and is the condition connecting the interventional model to qualitative probabilistic influence [23]. A local derivative at one operating point is insufficient by itself. Strict growth in the probability of an increased-value phenotype additionally requires positive probability mass to cross its reference threshold.

### S11 Derivative-sign abstraction

The structural causal model semantics is primary. The derivative-sign abstraction permits inference when complete functions, parameters, and distributions are unavailable. It does not change what the variables or typed relations mean. For an admissible model $M \in \mathfrak{M}(\mathcal{K}, \Pi)$, define

$$r^M_{k\to i}(u) = \left.\frac{\partial x_i^{*,\mathrm{do}(x_k=\theta)}(u)}{\partial\theta}\right|_{\theta=\theta_0}.$$

The ideal abstraction is

$$Q_{\mathcal{K}}(k,i) = \begin{cases} \texttt{+}, & r^M_{k\to i}(u) > 0 \text{ for all admissible } M \text{ and almost every } u, \\ \texttt{-}, & r^M_{k\to i}(u) < 0 \text{ for all admissible } M \text{ and almost every } u, \\ \texttt{0}, & r^M_{k\to i}(u) = 0 \text{ for all admissible } M \text{ and almost every } u, \\ \texttt{?}, & \text{otherwise.} \end{cases} \tag{S31}$$

The current first-order interface focuses on reached variables and reports +, -, ?; formally, an unaffected response 0 and an undetermined response ? are distinct.

Applying the sign map to the SCM derivatives gives the direct edge abstraction

$$\sigma_{ji} = \operatorname{sign}\left(\frac{\partial F_i}{\partial x_j}\right),$$

and the mixed-derivative abstraction is

$$\eta_{m;ji} = \operatorname{sign}\left(\frac{\partial^2 F_i}{\partial x_m\,\partial x_j}\right)$$

and $\lambda_{m;ji} = \sigma_{ji}\eta_{m;ji}$ states whether the magnitude of a sign-stable base influence is strengthened or weakened. The total second-order abstraction applies the same universal rule to Eq. (S28). For first-order computation, the derivative constraints associated with $E, \mathcal{P}, \mathcal{Q}$ form the signed dependency graph $G_\partial = (V, E_\partial)$. Constitutive constraints in $\mathcal{C}$ are applied separately, and modulations in $\mathcal{H}$ remain second-order constraints. The PhysioMap solver is derived from the SCM through this abstraction and is sound when every committed output agrees with the abstraction it implements. The reported experiments use its first-order query on $G_\partial$; the PhysioMap solver does not compute $\eta$, $\lambda$, quantitative values, or population orderings. Algorithmic failure to prove a first-order sign is conservative incompleteness; semantic ? means that admissible quantitative models genuinely disagree.

### S12 PhysioMap solver: algebra, proof, and algorithm

#### S12.1 Sign algebra

The following algebra operates only on the derivative-sign abstraction; it is not the semantics of the PhysioMap variables. Let the abstract sign domain be $\mathbb{S} = \{+, -, 0, ?\}$. A sign $s$ abstracts a real number $z$, written $s \triangleright z$, when + means $z > 0$, - means $z < 0$, 0 means $z = 0$, and ? admits every real value of the derivative or response being abstracted. Products along a path use $\otimes$; parallel contributions use $\oplus$:

| $\otimes$ | + | - | 0 | ? |
|---|---|---|---|---|
| + | + | - | 0 | ? |
| - | - | + | 0 | ? |
| 0 | 0 | 0 | 0 | 0 |
| ? | ? | ? | 0 | ? |

| $\oplus$ | + | - | 0 | ? |
|---|---|---|---|---|
| + | + | ? | + | ? |
| - | ? | - | - | ? |
| 0 | + | - | 0 | ? |
| ? | ? | ? | ? | ? |

.

Finite case analysis shows that these operations soundly abstract real multiplication and addition. In particular, opposing non-zero contributions return ? because their magnitudes are absent.

#### S12.2 Exact strongly connected components

Fix a strongly connected component $S$ with $m$ variables after intervention surgery. Let $J_S$ be its signed Jacobian, with off-diagonal entries given by causal-edge signs and negative diagonal entries representing local self-regulation. Let $b_S$ be the signed external forcing assembled from previously solved components. A real matrix is sign-consistent with $J_S$ when every entry has a sign admitted by the corresponding abstract entry.

**Comparative-static identity.** Consider a continuously differentiable realized model. Having solved the predecessor components, let $x^*_{\mathrm{up}}(\theta)$ collect their differentiable equilibrium responses. After substituting these responses into the equations for $S$, write

$$H_S(x_S,\theta) = F_S\bigl(x_S, x^*_{\mathrm{up}}(\theta), \theta\bigr) = 0.$$

At a reference intervention value $\theta_0$, define

$$J' = \left.\frac{\partial H_S}{\partial x_S}\right|_{(x^*_S,\theta_0)}, \qquad b' = \left.\frac{\partial H_S}{\partial \theta}\right|_{(x^*_S,\theta_0)}.$$

The vector $b'$ includes both direct forcing and forcing propagated from the previously solved components; $b_S$ is its sign abstraction. By assumption, $J'$ is Hurwitz and therefore nonsingular.

The implicit function theorem gives a locally unique differentiable equilibrium response $h_S$ satisfying $H_S(h_S(\theta),\theta) = 0$. Differentiation at $\theta_0$ gives

$$0 = J' h'_S(\theta_0) + b'.$$

With $z = h'_S(\theta_0) = dx^*_S/d\theta$, this is the linear system $J'z = -b'$. Let $J'^{(i)}$ replace column $i$ of $J'$ by $-b'$. Cramer's rule then gives

$$\frac{dx^*_i}{d\theta} = z_i = \frac{\det(J'^{(i)})}{\det(J')}. \tag{S32}$$

This proves the quantitative comparative-static identity used in the main paper.

**Selective-soundness statement.** Assume that the equilibrium is locally unique and that every consistent realized Jacobian $J'$ considered for the claim is Hurwitz. Let $J_S^{(i)}$ be $J_S$ with column $i$ replaced by $-b_S$. If the sign-only determinant $\mathrm{sdet}(J_S^{(i)})$ is `+` or `-`, then

$$\mathrm{sign}\!\left(\frac{dx^*_i}{d\theta}\right) = \mathrm{sdet}(J_S^{(i)})\,(-1)^m$$

for every sign-consistent stable realization and every sign-consistent forcing. If the signed determinant is `?`, the exact component algorithm abstains.

**Proof.** The signed determinant uses the same Laplace expansion as the real determinant, replacing multiplication and addition with $\otimes$ and $\oplus$. Induction on matrix dimension, together with soundness of the two abstract operations, shows that a definite $\mathrm{sdet}(A)$ equals the sign of $\det(A')$ for every sign-consistent real $A'$. A real Hurwitz $m \times m$ matrix has $\mathrm{sign}(\det J') = (-1)^m$: each real eigenvalue is negative, while each complex-conjugate pair contributes a positive product. Equation (S32) gives the response as the determinant ratio. Substitution of the numerator and denominator signs yields the stated result. Processing the condensation graph in topological order extends the result compositionally because upstream forcing is combined only with the sound sign operations. The combinatorial determinant argument is machine-checked in the Lean 4 development under `supplement/lean`; the eigenvalue statement is isolated there as the analytic assumption.

The theorem assumes a Hurwitz Jacobian and negative local self-regulation; PhysioMap does not infer either from clinical data. It applies to the exact component algorithm for every component it processes. The conservative fixed-point algorithm for larger components does not use the determinant denominator and may return `?` even when additional stability inequalities would determine a sign. Consequently, a committed exact-component sign has the universal guarantee above, while abstention may reflect either semantic indeterminacy or conservative algorithmic incompleteness.

**Exact component-algorithm cutoff.** The configurable component-size limit for the exact component algorithm was 16 traits in all reported analyses. For an $m$-variable component, the memoized determinant expansion has at most $2^m$ column-subset states for each numerator. Therefore, the cutoff is a computational resource guard, not part of the SCM semantics. The chosen value limits each expansion to at most $2^{16} = 65{,}536$ cached subset states.

We did not select 16 using the phenotype results. After every intervention used in the forward and inverse evaluations, non-singleton causal components contained either at most 10 variables or at least 188 variables; their observed sizes were $2, 3, 4, 8, 10, 188, 207, 210, 211, 212$, and 213. Consequently, every cutoff from 10 to 187 assigns exactly the same components to the two algorithms and produces the same reported predictions and ranks. The chosen value 16 lies within this invariance interval and is not scientifically distinguished.

##### S12.3 First-order inference algorithm

For an intervention $a$, the PhysioMap solver:

1. derives the signed dependency graph $G_\partial$ from the derivative constraints on authored causal influences and from the compatible first-order projections of production relations and quantitative identities;
2. replaces every intervened mechanism and removes its incoming signed dependencies;
3. computes strongly connected components and their condensation graph;
4. processes reachable components in topological order, using sign algebra for singletons, the exact component algorithm for components of size at most the configured limit, and the conservative fixed-point algorithm for larger components; and
5. applies constitutive determination between components without turning those constraints into causal edges.

All result traces identify the canonical content objects and intervention target. The PhysioMap solver reads relation type, derivative sign, and context, but it does not instantiate the quantitative random variables or functions and does not read the linked evidence and provenance metadata.

In release v1.1.1, the signed dependency graph contains 2,269 unique signed dependencies from 2,270 authored causal influences, 84 from production relations, and 15 from arguments to the results of 9 quantitative expressions. The PhysioMap solver applies 4 constitutive constraints separately. The PhysioMap solver does not read the 19 modulation relations. A separate qualitative query combines each modulation sign with a determinate first-order response of its modulator to report the direction in which the signed causal slope changes; it abstains when the required signs conflict or remain unresolved. The prediction and abduction evaluations did not score this second-order output.

Release v1.1.1 contains some exact quantitative identities and derivative constraints, but it does not contain complete numerical mechanisms, parameters, or exogenous distributions. Whole-map numerical or population inference therefore requires additional scientific content rather than activation of an unused solver option.

#### S13 Intended conditional-independence semantics

In a population realization with mutually independent exogenous variables and the appropriate unique-solvability conditions for the cyclic SCM, the observational distribution obeys the $\sigma$-Markov property [2, 4]. Conditional independences are therefore read by $\sigma$-separation. An exact target in $\Phi$ is a deterministic node: conditioning on all of its defining arguments fixes it and adds it to deterministic closure [5]. A modulation also creates ordinary parenthood $m \to i$ when the target mechanism depends on $m$; omitting that dependence would give incorrect separation claims.

### S14 Summary of the intended coupled semantics

An intended population model of PhysioMap is the coupled structure

$$\mathfrak{C} = (\mathcal{I}, \Omega, \Sigma, P, \{T_i, D_i, \rho_i, \nu_i, X_i\}_{i\in V}, E, \mathcal{H}, \{F_i\}, \Phi)$$

such that:

1. $\mathcal{I} \models \mathcal{K}$;
2. all $X_i$ are measurable on the same population space and satisfy $X_i = \nu_i \circ \rho_i$;
3. entailed class relations constrain phenotype events as in Eq. (S6);
4. every matched projection rule satisfies its quantitative conclusion;
5. the $F_i$ have the projected parent graph and functional constraints;
6. equations in $\Phi$ hold almost surely;
7. modulation hyperedges satisfy Eqs. (S22)–(S24);
8. the allowed observational and interventional regimes have the required solutions.

This specification leaves the standard OWL semantics unchanged. Its coupling to probability occurs through realization, valuation, and entailed projection rules. The projection has structural causal model semantics even when it specifies a class rather than a single parameterized model. Signs, abstention, and synergy labels are derived abstractions of the first- and second-order derivatives of those quantitative models. Release v1.1.1 materializes the ontology, typed manifest, derivative constraints, and PhysioMap solver, but not a complete member of this population model class.

### S15 Computational reproduction

#### S15.1 Construction provenance and source inventory

The released SCM and release manifest enumerate the complete list of compiled import files and their checksums. These files cover direct author curation, curated database imports, quantitative-model extraction, and textbook extraction. Every causal influence and modulation relation retains relation-level evidence text and a mechanism statement; other typed content objects preserve these annotations when they are available. Because older entries can have several sources and their acquisition route was not recorded uniformly, the release does not report mutually exclusive contribution counts for manual, database, model, and language-model acquisition.

The quantitative cardiovascular source set is enumerated in `benchmarks/guyton/SOURCES.md`. It contains the following 17 Guyton control modules downloaded through the BioModels REST API: `MODEL0911342562`, `MODEL0911376350`, `MODEL0911309080`, `MODEL0911272039`, `MODEL0911270005`,

MODEL0909931851, MODEL0911270006, MODEL0912160001, MODEL0912160000, MODEL0911169699, MODEL0911202318, MODEL0910896131, MODEL0910928451, MODEL0911231713, MODEL0910846879, MODEL0911091440, and MODEL0911047946. The CellML workspaces guyton_circulatory_dynamics_2008, guyton_kidney_2008, and guyton_2008 supply the circulatory, renal, and full-model integrators; BioModels MODEL2202160001 is a structural cross-check. For each imported influence, its provenance identifies the source equation or component, and the sign is the sign of the target equation's partial derivative with respect to the source variable. Curated database inputs include the versioned SIGNOR metabolism and AOP-Wiki import files under benchmarks/human/curated/.

The historical language-model batch used OpenStax *Anatomy & Physiology 2e* (169 sections), Wong's *Cells: Molecules and Mechanisms* (29 sections), and Jakubowski and Flatt's *Fundamentals of Biochemistry*, volume I (60 sections). Licenses, source URLs, and local filenames are recorded in resources/textbooks/README.md; section manifests and extraction summaries are under benchmarks/results/. Direct author curation and later literature corrections are retained in the compiled input files and the per-relation evidence annotations. This inventory identifies all machine inputs and external collections used by the release; the historical route of every individual content item cannot be reconstructed where that metadata was never recorded.

##### S15.2 Historical language-model proposal batch

The expansion batch summarized in the main paper is documented by the source inventory in resources/textbooks/README.md, the public run summary in benchmarks/results/textbook_extraction.md, and the integrated fragment benchmarks/human/systems/textbook_extracted.yaml. The run summary reports Claude Sonnet extraction and separate adversarial verification with Claude Opus. Three open-licensed textbooks were divided into 258 sections. The batch made 362 model calls and processed approximately 10.8 million tokens. It proposed 111 non-duplicate nodes and 228 distinct signed relations; semantic and scale filtering retained 111 nodes and 87 causal influences for integration.

The public release does not contain the original orchestration implementation, immutable model snapshot identifiers, sampling parameters, complete invocation metadata, section-level structured outputs, or internal reconciliation transcripts. Consequently, the integrated fragment and its retained relation-level evidence are auditable, but the calls and reported processing totals cannot be reconstructed or replayed exactly from the public archive. The retained proposals passed a separate model review and automated schema, ontology-identifier, relation-type, and regression checks. Most did not receive independent edge-by-edge human source review; their evidence classes must not be interpreted as human approval.

##### S15.3 Stratified expert review of released content

On 2026-07-28, a physiologist (P.N.S.) received a fixed-seed sample of 83 relation assertions from the v1.1.1 SCM export. The sampling frame contained 2,270 causal influences, 85 production relations, 19 modulations, four constitution constraints, and nine quantitative identities. The sample included 50 causal influences stratified by evidence class, simple random samples of 10 production relations and 10 modulations, and all constitution constraints and quantitative identities. The causal allocation was approximately proportional, subject to at least five sampled assertions from every evidence class containing at least five assertions. The seed was 20260728. Because the relation types were sampled at different rates, the unweighted totals describe the reviewed sample and are not a map-wide accuracy estimate.

Each workbook row showed the relation type and arguments, the sign where applicable, evidence class, mechanism, and evidence text. P.N.S. assessed the relation content against his physiological knowledge. The review was therefore not blinded to the supporting information. We interpret `TRUE` as accepted, `FALSE?` as flagged for further review or investigation because the assertion may be false or was not immediately verifiable, and `FALSE` as rejected as false. A flagged assertion is not a rejected assertion.

Table S1: Stratified expert review of released PhysioMap content. Flagged relations require further review or investigation; rejected relations were judged false.

| relation type | accepted | flagged | rejected | reviewed |
|---|---|---|---|---|
| Causal influence | 42 | 6 | 2 | 50 |
| Production | 8 | 2 | 0 | 10 |
| Constitution | 4 | 0 | 0 | 4 |
| Quantitative identity | 8 | 1 | 0 | 9 |
| Modulation | 7 | 3 | 0 | 10 |
| Total | 69 | 12 | 2 | 83 |

The returned workbook contains a verdict for every sampled assertion and comments on 39. The importer verified that all 83 identifiers and the relation, sign, evidence-class, mechanism, and evidence cells were unchanged from the sent workbook. Checksums and archive metadata for the sampling frame, sent workbook, and returned workbook are listed in the machine-readable expert-review summary. The versioned review archive contains the exact sampling frame, sent and returned workbooks, complete row-level judgments in JSON and TSV, checksums, and the sampling and import scripts.

The comments raise specific curation questions about variable scope, conflated processes, causal directness and context, and operating-point-dependent modulation. These observations guide relation-level review but do not define additional outcome categories. The review has one expert, small type-specific samples, unequal sampling fractions, and visible supporting information. It therefore provides direct cross-type scrutiny without establishing a population estimate of correctness.

#### S15.4 Prediction and abduction reference inputs

The evaluation uses the tagged HPO release 2026-02-16. Checksum-pinned compressed copies of `hp.obo` and `genes_to_phenotype.txt` are versioned under `benchmarks/data/hpo-2026-02-16/`. The release gate verifies the compressed archives, expands them into a local cache, and verifies the expanded files before evaluation. Checksums for the frozen inputs are listed in the machine-readable evaluation manifest. The corresponding tagged assets are available from `https://github.com/obophenotype/human-phenotype-ontology/releases/tag/v2026-02-16`.

The candidate lesion specification contains 184 genes and 188 primary variable assignments. We mapped 163 of these genes by joining the approved gene symbol to a UniProt identifier in the HGNC export, joined UniProt to PR through PRO `promapping.txt`, and matched the resulting PR class to the bearer of an enzyme- or transporter-activity variable in PhysioMap. Loss-of-function disorders were represented as decreases in the matched activity. GLUD1 and PRPS1 activating disorders were represented as increases. We retained 21 manually curated mappings whose intervention variable and direction had been selected from the disease mechanism; these include cases represented by a hormone or a proximal physiological function rather than directly by the gene product. Four genes, ARSA, CTNS, KYNU, and PKLR, map to two primary variables. The complete gene-to-intervention list is the versioned lesion specification. HPOA uses the symbol G6PC1 where this specification uses G6PC, so the exact-symbol join matched 183 genes. Sixteen matched genes had only a direction for the intervened variable, which was not scored. The downstream reference directions before discrepancy exclusions therefore come from the remaining 167 genes. The single-lesion inverse benchmark excludes the four genes with two primary variables, which leaves 163 genes.

Let $A_g$, $\mathsf{HPO}$, and $M$ have the meanings defined in the main paper. For every direct HPOA annotation $h$, the construction tested $h$ and every superclass $a$ for which $\mathsf{HPO} \models h \sqsubseteq a$. A mapped superclass contributed its PhysioMap variable and PATO increased-quality or decreased-quality direction. For example, fasting hypoglycemia (HP:0003162) satisfies

$$\mathsf{HPO} \models \mathsf{FastingHypoglycemia} \sqsubseteq \mathsf{Hypoglycemia},$$

and the curated mapping of hypoglycemia contributes $(\mathsf{plasma\ glucose}, \mathtt{-})$. No subclass of an annotated term was considered. If this construction yielded both directions for the same gene and variable, that pair was discarded.

The lesion and directional term specifications are `benchmarks/hpo/gene_lesions_e1b.yaml` and `benchmarks/hpo/hpo_term_map.yaml`. The complete directional map is also available as a versioned HPO-to-PhysioMap specification. Propagation is blocked for HP:0025436 (elevated serum 11-deoxycortisol) and HP:6000516 (elevated circulating 21-deoxycortisol), because neither denotes cortisol despite its asserted ancestry. Before discrepancy exclusions, the reference contained 870 downstream pairs. The PhysioMap solver made 175 determinate predictions, of which 171 agreed with the HPO-derived direction and four disagreed. The four exclusions in `benchmarks/hpo/annotation_discordances.yaml` are LDHA–plasma lactate, ABCA1–plasma apolipoprotein A-II, LPL–LDL cholesterol, and LCAT–LDL cholesterol. The first three mapped directions conflict with the cited primary literature; the last is variable. These four exclusions were identified during discrepancy review and were not prespecified. The evaluation applies them uniformly to every method. Without them, the four PhysioMap solver predictions are counted as contradictions. The adjudicated set of 866 pairs is reconstructed deterministically from the two versioned specifications, the checksum-pinned HPO assets, and the documented exclusions. The complete row-level set is materialized as the versioned row-level evaluation table; each row gives the gene, primary intervention, PhysioMap variable, reference direction, direct HPO annotation, mapped directional HPO class, and lesion-mapping note.

For the gene-stratified abstention test, we restricted the reference to pairs on which the shortest-path rule returned a direction. For gene $g$, let $n_g$ be the number of these pairs, $a_g$ the number on which the PhysioMap solver abstained, $e_g$ the number of shortest-path errors, and $Z_g$ the number of those errors among the abstained pairs. Under the null hypothesis, conditional on these margins, the $a_g$ abstention labels are exchangeable among the $n_g$ shortest-path predictions within gene $g$. Every feasible $z$ then has probability

$$\Pr(Z_g = z \mid n_g, a_g, e_g) = \frac{\binom{e_g}{z}\binom{n_g - e_g}{a_g - z}}{\binom{n_g}{a_g}}. \tag{S33}$$

We convolved these gene-specific hypergeometric distributions under independent randomization across gene strata and computed the exact one-sided upper-tail probability of $Z = \sum_g Z_g$. This conditional randomization test does not assume that phenotype pairs from the same gene are independent, but it does assume within-gene exchangeability of the abstention labels under the null. The implementation and generated result are archived in repository revision 79ff2fb.

The leakage sensitivity analysis excludes ADA, ADA2, ADSL, AK1, AMPD1, APRT, DPYD, DPYS, ITPA, PRPS1, TYMP, UGT1A1, UMPS, UPB1, and XDH.

##### S15.5 Release and commands

Experiments use PhysioMap v1.1.1 with Python 3.11 or newer. The canonical input is `release/owl-scm/physiomap-scm.json`. Its checksum is listed in the machine-readable release checksum manifest. Projection rules are fixed by `projection/patterns.yaml`, and Python dependencies by `uv.lock`. Repository revision 79ff2fb retains the released v1.1.1 content and frozen inputs and adds the gene-stratified test. From that revision, the reported results and figures are regenerated by:

```
uv sync --frozen --extra analysis
uv run python scripts/bootstrap_release_inputs.py
uv run python scripts/build_hpo_observations.py
uv run python scripts/e1b_eval.py --leakage-sensitivity
uv run python scripts/e2_baseline.py
uv run --extra analysis python scripts/e2c_risk_coverage.py
uv run python scripts/e4_diagnose.py
uv run --extra analysis python scripts/e4b_diagnosis_baselines.py
uv run python scripts/e5_typed_layer_ablation.py
uv run --extra analysis python \
  scripts/generate_psb_rare_disease_figure.py
uv run python scripts/build_expert_gold_sample.py --seed 20260728
uv run python scripts/import_expert_gold_review.py --check
uv run python scripts/owl_scm_release_gate.py
```

The figure generator reads the evaluation outputs and rewrites the result macros used by the paper. Composition statistics and tables are generated from the canonical SCM by `scripts/generate_migration_statistics.py`. The typed-layer ablation rewrites `benchmarks/results/e5_typed_layer_ablation.json` and its Markdown summary. When supplied an explicit output path in a linked paper checkout, it also rewrites the corresponding LaTeX table. Because it repeats the inverse pool under four configurations, it should be run on a compute host. The final command rebuilds the authoritative projection and rejects stale checksums, ontology inputs, statistics, evaluation outputs, generated figures, behavioral baselines, or public-interface artifacts. Paper compilation can additionally be enforced from a linked paper checkout.

#### S16 Generated content characterization

All values and figures in this section are generated from the released OWL, SCM manifest, projection witnesses, locality-module manifest, and versioned behavioral reference. Release validation rejects stale generated artifacts.

Table S2: Sensitivity to cumulative first-order relation types. Forward coverage reports determinate calls among 866 adjudicated pairs; inverse results use the true lesion's best-tied rank.

| active relation types | determinate | coverage | top-3 | MRR |
|---|---|---|---|---|
| causal | 173 | 20.0% | 129/163 | 0.710 |
| causal + production | 172 | 19.9% | 127/163 | 0.707 |
| causal + production + quantitative | 171 | 19.7% | 127/163 | 0.707 |
| causal + production + quantitative + constitution | 171 | 19.7% | 127/163 | 0.707 |

Table S3: Trait completeness.

| Trait status | Count | Percent |
|---|---|---|
| Complete | 1568 | 92.3% |
| Incomplete primitive | 131 | 7.7% |

Table S4: Registered projection matches.

| Pattern | Unique entailments |
|---|---|
| causal-collection-v2 | 2269 |
| constitution-v1 | 4 |
| multiplicative-modulation-v2 | 19 |
| production-collection-v2 | 85 |
| ratio-v1 | 1 |

Table S5: Recovery of the migration-source model.

| Migration check | Recovered | Expected |
|---|---|---|
| Nodes | 1699 | 1699 |
| Contextual influences | 2270 | 2270 |
| Production relations | 85 | 85 |
| Constitutive edges | 4 | 4 |
| Quantitative definitions | 9 | 9 |
| Modulations | 19 | 19 |

Table S6: Validation outcomes.

| Release gate | Outcome |
|---|---|
| OWL 2 EL profile | passed |
| ELK consistency | passed |
| ELK classification | passed |
| Projection equivalence | passed |
| Derivative rules | 38 |
| Numerical realizations | 448 |

Table S7: Locality-module refresh performance.

| Source module | Axioms | Extraction (s) |
|---|---|---|
| CHEBI.obo | 27009 | 32.431 |
| CL.obo | 4776 | 6.841 |
| GO.obo | 5793 | 17.216 |
| PATO.obo | 681 | 4.998 |
| PR.obo | 38142 | 129.487 |
| UBERON.obo | 13260 | 9.568 |

Table S8: Physiology benchmark outcomes.

| Benchmark | Cases | Correct | Wrong | Ambiguous |
|---|---|---|---|---|
| drug_panel | 10 | 14 | 0 | 17 |
| guyton | 4 | 22 | 2 | 6 |
| human | 77 | 91 | 0 | 190 |
| human_multiscale | 2 | 5 | 0 | 2 |

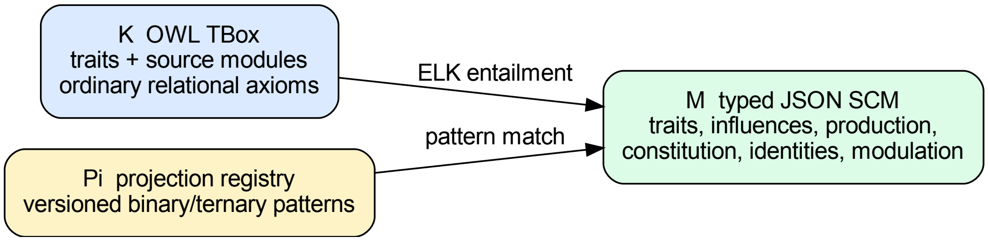


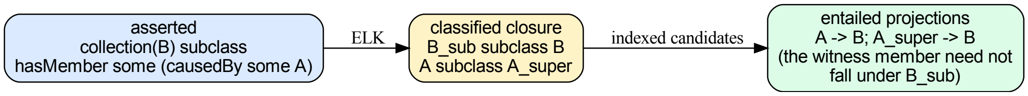


Figure S1: Projection to structural causal model semantics (top), and asserted-to-inferred projection (bottom).

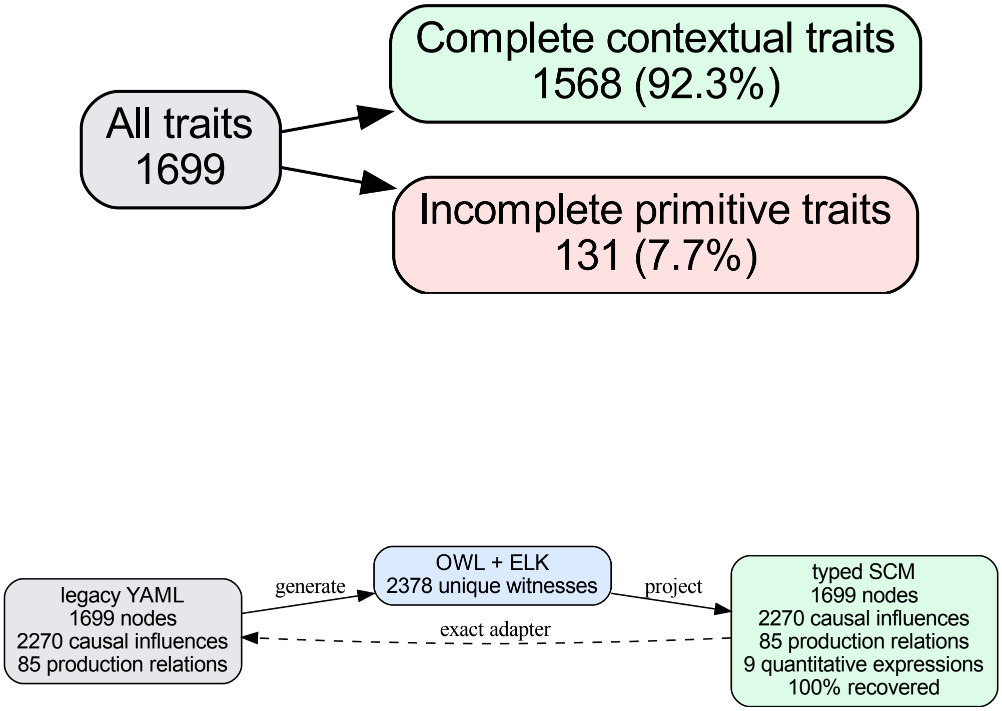


Figure S2: Trait completeness (top) and legacy-model recovery (bottom).

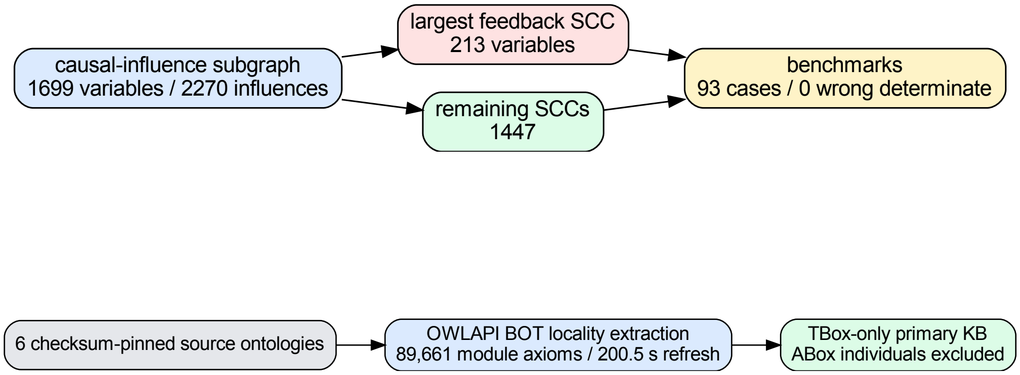


Figure S3: Causal-influence subgraph structure (top) and locality extraction (bottom).

#### S17 Generated construction-metadata characterization

This table characterizes the separate evidence-review state used to construct and audit the causal content. Its categories do not enter the content projection or any reported application.

Table S9: Causal-evidence migration status.

| Causal-evidence status | Count |
|---|---|
| Controlled | 2117 |
| Legacy unclassified | 153 |
| Frozen review baseline | 621 |
| Human-approved resolutions | 470 |
| Open review items | 151 |

#### S18 Relation to prior work

The EQ definitions follow the phenotype-ontology tradition [6, 7]. The separation of an OWL knowledge base from a projected causal subset extends the causal knowledge graph semantics of Toonsi et al. [22], while the realization and valuation maps generalize their binary population events to real-valued traits. The reasoner-first projection follows Onto2Graph [18]. The dynamical and equilibrium semantics follows work connecting interventions on differential equations to cyclic SCMs [2, 12, 19]. The measurement rules use classical extensive and ratio measurement theory [11, 17, 20]; the interventionally consistent constitutive maps are causal abstractions in the sense of Beckers and Halpern [1]. PhysioMap combines these components in an OWL-grounded representation in which classified trait and relational patterns constrain one joint quantitative causal and probabilistic model, including explicit second-order modulation. The signed dependency graph and the PhysioMap solver are derived as a sound first-order abstraction of that model class.
